# A feedforward loop between STAT1 and YAP1 stimulates lipid biosynthesis, accelerates tumor growth, and promotes chemotherapy resistance in mutant KRAS colorectal cancer

**DOI:** 10.1101/2024.10.03.616541

**Authors:** Shuo Wang, Shiqi Diao, Hyungdong Kim, Jia Yi Zou, Ke Ke Li, Antonis E. Koromilas

**Affiliations:** Lady Davis Institute for Medical Research, McGill University, Sir Mortimer B. Davis-Jewish General Hospital, Montreal, Quebec H3T 1E2, Canada; Graduate Program in Clinical and Translational Research, Faculty of Medicine, McGill University, Montreal, Quebec H4A 3J1, Canada; Department of Microbiology and Immunology, School of Biological Sciences, McGill University, Montreal, Quebec H3A 2B4, Canada; Gerald Bronfman Department of Oncology, Faculty of Medicine, McGill University, Montreal, Quebec H4A 3T2, Canada

**Keywords:** Signal transducer and activator of transcription 1, Ki-ras2 Kirsten rat sarcoma viral oncogene homolog, sterol regulatory element binding proteins, mevalonate pathway, Yes-associated protein, tumorigenesis

## Abstract

In tumorous conditions, the transcription factor STAT1, traditionally recognized for its anti-tumor role in immunology, exhibits pro-survival characteristics, though the underlying mechanisms remain unclear. Investigating STAT1’s function in isogenic colorectal tumor cells with wild-type or mutant KRAS, we found that STAT1 specifically promotes the survival and proliferation of cells with mutant KRAS. Through gene expression profiling, we discovered a previously unknown role of STAT1 in upregulating sterol and lipid biosynthetic genes specifically in mutant KRAS cells. This effect is driven by STAT1’s phosphorylation at serine 727 and its cooperation with STAT3 and STAT5 for the transcriptional upregulation of sterol regulatory element-binding proteins (SREBP) 1 and 2, which boost *de novo* sterol and lipid biosynthesis. In mutant KRAS cells, STAT1 amplifies the mevalonate pathway, maintaining its serine 727 phosphorylation through RHO GTPase signaling and establishing a positive feedback loop through the transcription factors YAP1 and TEAD4, further driving lipid biosynthesis and tumor growth. Through xenograft tumor assays in mice, we discovered that the STAT1-YAP1 axis plays a role in mutant KRAS tumor cells’ resistance to mevalonate pathway inhibitors, which can be overcome by pharmacologically targeting the YAP1-TEAD interaction. Additionally, the STAT1-YAP1 arm is essential for the intrinsic resistance to EGFR-targeting therapy in the mutant KRAS colon cancer cells. These findings indicate that the STAT1-YAP1 pathway plays a significant role in therapy resistance and presents a potential therapeutic target in mutant KRAS colorectal cancer.

## INTRODUCTION

The RAS family, including HRAS, NRAS, and KRAS, comprises small membrane bound GTPases of approximately 189 amino acids. These proteins are frequently mutated in about 30% of human cancers ^1^. Somatic gain-of-function *RAS* mutations typically involve a single amino acid substitution, primarily at G12, G13, or Q61, resulting in conformational changes that favor the protein’s presence in an active GTP-bound form ^1^.

Among the RAS isoforms, Ki-ras2 Kirsten rat sarcoma viral oncogene homolog (KRAS) is the predominant one mutated in three of the most lethal cancers globally, namely colorectal, lung, and pancreatic cancers ^2^. Mutant KRAS accelerates proliferation and establishes crosstalk between tumors and stromal cells in the tumor microenvironment, overcoming challenges such as low nutrient availability and immune-mediated anti-tumor responses ^3^. Additionally, mutant KRAS alters key metabolic processes, including elevating glucose, lipids, glutamine, and glucose uptake and consumption, to support biosynthetic pathways and enhance cellular redox potential^3^. Tumors harboring *KRAS* mutations exhibit dependence on these metabolic adaptations, suggesting potential novel avenues for cancer therapy^3^.

The signal transducer and activator of transcription 1 (STAT1) plays a crucial role in innate immunity, safeguarding the host against infections caused by viruses and other pathogens ^4^. Its primary functions are well-defined downstream of type I (α/β) and type II (γ) interferon (IFN) receptors, where it actively participates in the expression of anti-proliferative, anti-viral, and immune regulatory genes ^5^. In the realm of cancer research, STAT1 demonstrates anti-tumor effects in mice treated with carcinogens, primarily through the stimulation of anti-tumor immunity^6^. STAT1 utilizes both tumor intrinsic and immune-mediated mechanisms to suppress the growth of breast cancers in mouse models of the disease^6^. However, STAT1 can also exhibit oncogenic effects by disrupting immune surveillance mechanisms against certain types of blood tumors^6^. Despite its context-dependent role in cancer formation, STAT1 renders solid cancers resistant to chemotherapy and radiotherapy via the stimulation of IFN-inducible genes that confer resistance to DNA damage ^6,7^. Besides its role in gene transcription, STAT1 promotes cell survival by activating the phosphoinositide 3-kinase (PI3K)-mammalian target of rapamycin (mTOR) pathway and enhancing the mRNA translation of anti-apoptotic genes ^8-10^.

YES-associated protein 1 (YAP1) is hyperactivated in numerous human cancers and demonstrates tumorigenicity in mouse cancer models ^11^. YAP1 is a crucial target for inactivation by the HIPPO tumor suppressor pathway, which is modulated by diverse signals, including oncogenic and metabolic determinants ^12^. Mutant KRAS counteracts the HIPPO pathway, amplifying the stabilization, nuclear localization, and gene transactivation capacity of YAP1 ^13,14^. Mutant KRAS promotes the nuclear localization and gene transactivation capacity of YAP1 through both HIPPO-dependent and independent pathways ^13-15^. YAP1 displays a pro-survival role in colorectal, lung, and pancreatic cancers with *KRAS* mutations ^16-19^.

Tumors often exploit alterations in metabolic pathways to generate intermediates necessary for rapid proliferation ^20^. One such pivotal pathway in tumor proliferation is *de novo* lipogenesis ^20^. While normal cells typically rely on lipid uptake from circulation for their metabolic needs, tumors respond to the heightened demand for membrane biogenesis during proliferation by activating *de novo* sterol and lipid biosynthetic pathways ^21^. Sterol regulatory element-binding proteins (SREBPs) 1 and 2, transmembrane proteins located in the endoplasmic reticulum, play a crucial role in this process ^22^. Depletion of sterols or unsaturated fatty acids triggers the trafficking of SREBPs to the Golgi apparatus, where proteolytic cleavage activates them, allowing entry into the nucleus to upregulate the expression of cholesterol and lipid biosynthetic genes ^22,23^. There are three isoforms of SREBPs; the alternatively spliced forms SREBP-1a and SREBP-1c, with the latter being dominant in cultured cells, activate transcription of genes involved in fatty acid synthesis, while SREBP-2 upregulates enzymes required for cholesterol biosynthesis.

Mutant KRAS elevates *de novo* lipid biosynthesis by stimulating sterol regulatory element-binding proteins (SREBPs) and fatty acid synthase (FASN) activity ^24-26^. However, the specific pathways involved in *de novo* lipogenesis in mutant KRAS cancer cells remain partially understood. STAT1 is known for its role in cholesterol metabolism through the transcriptional upregulation of 25-hydroxycholesterol in macrophages as part of an anti-viral response induced by interferons (IFNs) ^27^. Conversely, YAP1 is involved in upregulating *de novo* lipogenesis in non-tumorigenic tissue and diabetic mouse models ^28,29^. However, it remains unknown whether STAT1 or YAP1 are implicated in lipogenic programs of tumor cells.

The upregulation of STAT1 in human colon adenocarcinomas, compared to normal epithelial cells ^30^, suggests a potential role for STAT1 in promoting colorectal cancer (CRC) growth. Using isogenic CRC cells with either wild-type or mutant KRAS, we demonstrate that STAT1 specifically promotes the survival and growth of tumor cells with mutant KRAS. The pro-survival function of STAT1 is intricately linked to its ability to enhance the expression of SREBPs and downstream sterol and lipid biosynthetic genes in mutant KRAS cells. Our findings reveal that the activation of the mevalonate pathway by STAT1 leads to increased nuclear localization and enhanced transcriptional function of YAP1. Subsequently, activated YAP1 forms a complex with the transcription factor TEAD4, thereby increasing the expression of SREBPs and lipogenic gene expression in mutant KRAS cells.

Our data underscore the establishment of a feedforward loop between STAT1 and YAP1, driving the upregulation of lipogenic pathways and conferring resistance to mevalonate pathway and EGFR inhibitors in mutant KRAS cells. This intricate interplay reveals a potential target for therapeutic intervention in the context of mutant KRAS-associated cancers.

## RESULTS

### STAT1 promotes the survival of colorectal tumor cells with mutant but not wild type KRAS

In our investigation, we utilized human CRC HCT116 and DLD1 cells expressing a *KRAS G13D* allele, along with their isogenic counterparts HK2-8 and DKO-1 cells, which harbor wild-type (WT) *KRAS* ^31^. The downregulation of STAT1 using siRNAs resulted in a significant decrease in the colony-forming efficacy of HCT116 and DLD1 cells, whereas the colony-forming efficacy of HK2-8 and DKO-1 cells remained unaffected (Fig. 1a; Supplementary Fig. 1a). Immunoblotting validated the reduction of STAT1 in these isogenic colon cancer cells (Fig. 1b; Supplementary Fig. 1b). We tested the specificity of STAT1 downregulation by verifying that expression of STAT3 and STAT5 upon treatment with STAT1 siRNAs remained unimpaired (Fig. 1b). These data collectively emphasize the pro-survival function of STAT1 specifically in tumor cells with mutant KRAS, with no discernible impact on cells with wild-type KRAS.

**Fig. 1.**
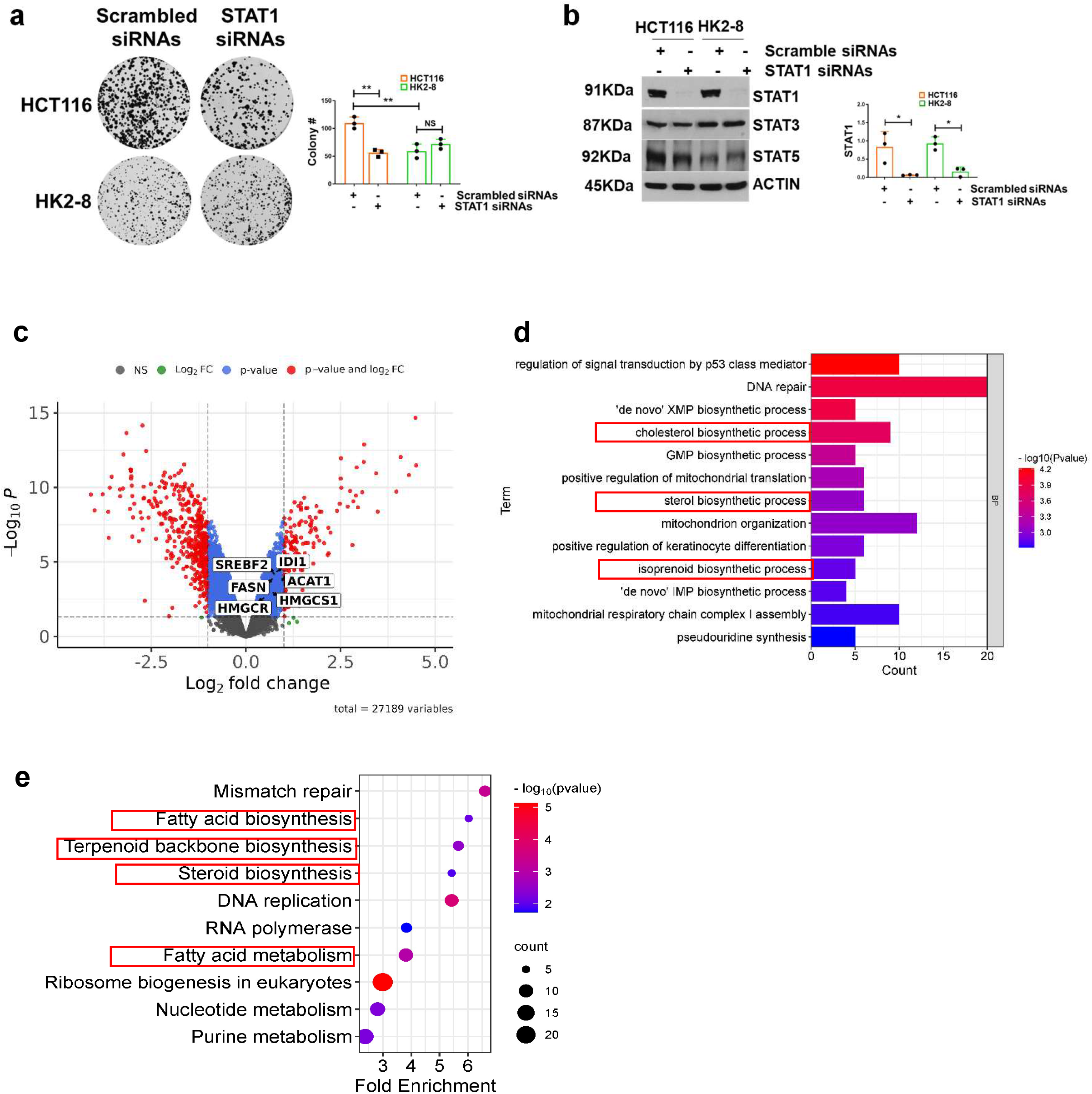
STAT1 is a pro-survival factor and promotes the expression of sterol and lipid biosynthetic genes in mutant KRAS cells. (**a**) Evaluation of colony formation efficacy of HCT116 and HK2-8 cells after treatments with either scramble siRNAs or STAT1 siRNAs. The graphs represent data obtained from 3 biological replicates each of which included 3 technical replicates and represent ±SEM (*\**) *P* < 0.05 (*\*\**) *P* < 0.01;(*t*-test), NS=non-significant. **(b)** Immunoblotting of indicated protein from extracts of siRNA-treated cells. Quantifications show the relative intensity of STAT1 normalized to ACTIN from 3 biological replicates and represent ±SEM (*\**) *P* < 0.05;(*t*-test). (**c**) Volcano Diagram illustrating the number of genes differentially expressed in STAT1-replete (WT; control) compared to STAT1^-/-^ HCT116 cells. Essential genes of sterol and lipid biosynthetic pathways are indicated. (**d**) Bar plot of biological processes (BP) from gene ontology (ON) significantly enriched in STAT1-dependent genes in HCT116 cells. (**e**) Bot plot of Kyoto Encyclopedia of Genes and Genomes (KEGG) pathways under the control of STAT1 in HCT116 cells.

### STAT1 upregulates SREBPs and lipid biosynthetic genes in mutant KRAS tumor cells

To explore the mechanisms underlying STAT1 function, we conducted a comparative analysis of gene expression profiles between proficient (STAT1^+/+^) and STAT1-deficient (STAT1^-/-^ via CRISPR) HCT116 cells. Verification of STAT1 deletion was accomplished through immunoblotting, both before and after treatment with IFNs, known to induce STAT1 expression (Supplementary Fig. 1c). Our analysis revealed 1922 genes that were upregulated, and 2058 genes downregulated by STAT1 (Fig. 1c). Subsequent gene ontology (GO) enrichment and pathway assessments unveiled the involvement of STAT1-regulated genes in diverse biosynthetic processes, with a prominent emphasis on the upregulation of steroid and lipid biosynthetic pathways (Fig. 2 d, e).

**Fig. 2.**
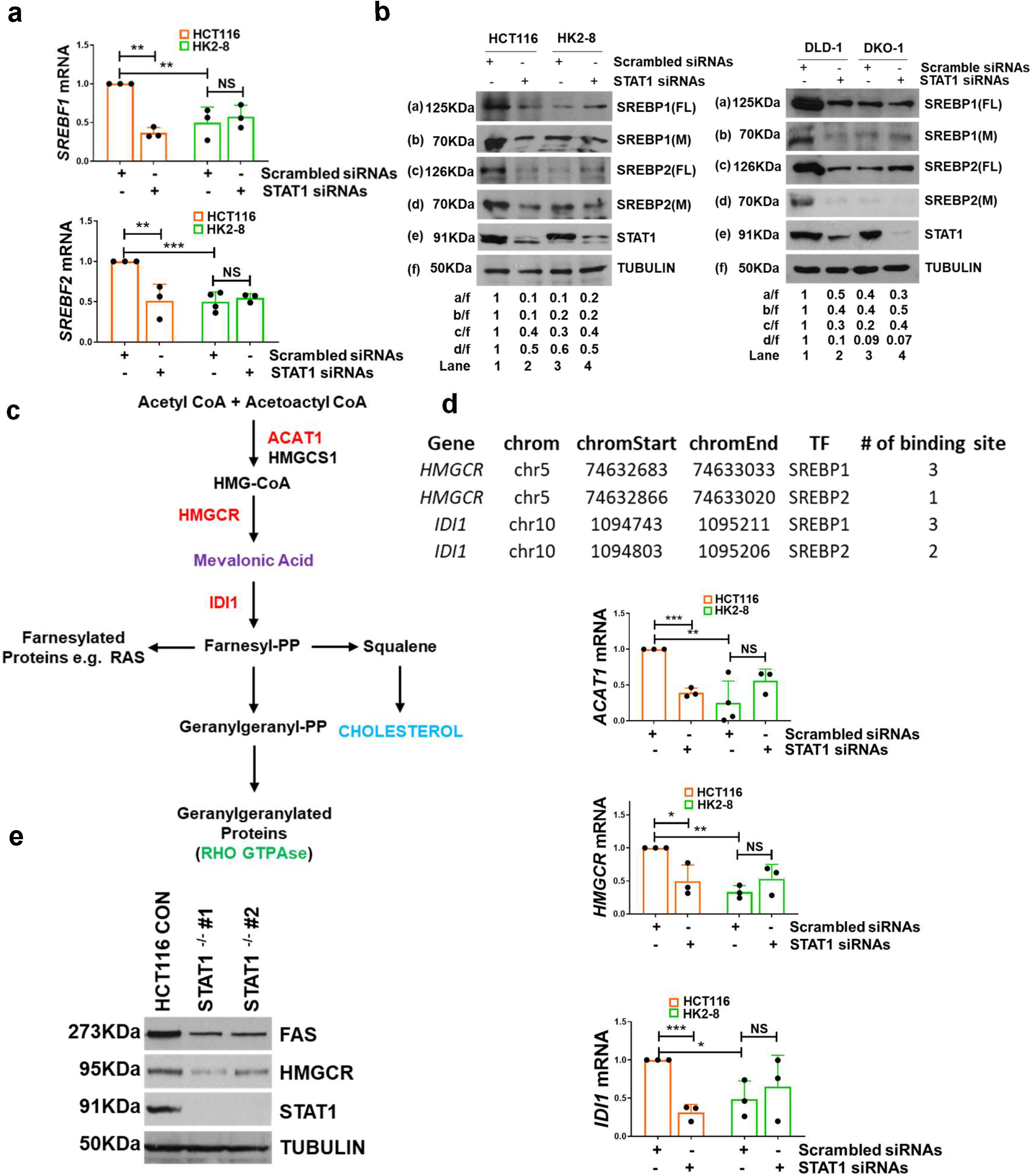
STAT1 upregulates SREBPs and mevalonate pathway genes in mutant KRAS cells. (**a**) Detection of *SREBF* mRNAs in colon cancer cells with intact or impaired STAT1. *SREBF 1* and *2* mRNA levels were normalized to ACTIN and TUBULIN mRNAs used as internal controls. Data obtained from 3 biological replicates each of which contained 3 technical replicates and represent ±SEM (*\*\**) *P* < 0.01; (*t*-test), NS=non-significant (**b**) Immunoblotting for SREBP1 and 2 by immunoblotting in colon cancer cells with intact or downregulated STAT1. Quantifications show the relative intensity of proteins normalized to TUBULIN. FL, full length; M, mature form of SREBPs. (**c**) Schematic representation of the mevalonate pathway. Genes in red color are SREBP-dependent genes. (**d**) ChIP-seq data from ENCODE (UCSC data base) indicating the binding of SREBP1 and 2 to transcriptional regulatory regions of mevalonate pathway genes. Graphs show the detection of *ACAT1, HMGCR* and *IDI1* mRNAs by qPCR in cells treated with scrambled or STAT1 siRNAs. Gene expression was normalized to ACTIN and TUBULIN mRNAs used as internal controls. Data were obtained from 3 independent experiments performed in triplicates and represent ±SEM (*\**) *P* < 0.05; (*\*\**) *P* < 0.01; (***) *P* < 0.001 (*t*-test). (**c**) Immunoblotting of HCT116 protein extracts replete (control) or deplete (^-/-^) for STAT1 by CRISPR (cell line #1 and #2). Detection of STAT1 and rate-limiting enzymes of sterol and lipid biosynthetic pathway HMGCR and FAS, respectively.

The analysis of gene expression data revealed STAT1’s involvement in the upregulation of *SREBF-1* and *SREBF-2* mRNAs, responsible for encoding SREBP-1 and SREBP-2, respectively (Fig. 1c). These transcription factors act as master regulators for cholesterol and lipid biosynthetic genes ^32,33^. To confirm this, we employed STAT1 siRNAs to downregulate STAT1 and observed a reduction in *SREBF-1* and *2* mRNA levels in HCT116 cells carrying the *KRAS G13D*, but not in isogenic HK2-8 cells with wild-type *KRAS* (Fig. 2a). Further, treatment with STAT1 siRNAs resulted in decreased expression of both the full-length (FL) 125 kDa and mature (M) 70 kDa forms of SREBP-1 and 2 in HCT116 and DLD-1 cells harboring mutant KRAS, while no such effect was observed in isogenic HK2-8 and DKO-1 cells with wild-type KRAS (Fig. 2b). These findings support the conclusion that the STAT1-mediated upregulation of SREBPs occurs specifically in cells with mutant KRAS.

Within the genes upregulated by STAT1, acetyl-CoA acetyltransferase (*ACAT1*), 3-hydroxy-3-methylglutaryl-CoA reductase (*HMGCR*), and isopentenyl-diphosphate delta-isomerase 1 (*IDI1*) encode pivotal enzymes in the mevalonate pathway, a critical player in the synthesis of cholesterol and isoprenoid metabolites (Fig. 2c) ^34^. Analysis of the Encyclopedia of DNA Elements (ENCODE) revealed the presence of SREBP1 and 2 binding sites in the regulatory regions of these genes, implying an indirect role of STAT1 in their expression through the upregulation of SREBPs (Fig. 2d). Indeed, STAT1 downregulation led to a reduction in the expression of *ACAT1, HMGCR*, and *IDI1* mRNAs in HCT116 cells, while no such effect was observed in isogenic HK2-8 cells (Fig. 2d). Immunoblotting of HCT116 cells further confirmed that the deletion of STAT1 by CRISPR reduced the expression of SREBP-dependent gene products, such as HMGCR, the rate-limiting enzyme of the mevalonate pathway^34^, and fatty acid synthase (FAS), the rate-limiting enzyme of fatty acid biosynthesis in colon cancers ^35^ (Fig. 2e). These findings suggest that STAT1, acting through SREBPs, fosters the expression of key enzymes in sterol and lipid biosynthetic pathways specifically in cells with mutant KRAS.

### STAT1 stimulates the transcription of SREBF1 and 2 genes specifically in mutant KRAS cells

SREBF1 and 2 are transcribed from distinct promoters ^36^. Analysis of the Encyclopedia of DNA Elements (ENCODE) database revealed two STAT1 binding sites in the *SREBF-1* gene and one in *SREBF-2* (Fig. 3a). Chromatin immunoprecipitation (ChIP) assays showed the binding of endogenous STAT1 to the regulatory regions of *SREBF1* and *2* in HCT116 cells compared to HK2-8 cells (Fig. 3b). As STAT1’s transcriptional activity is modulated by phosphorylation at tyrosine (Y) 701 and serine (S) 727 ^4^, we investigated if phosphorylation at these sites influences the transcription of *SREBF* genes.

**Fig. 3.**
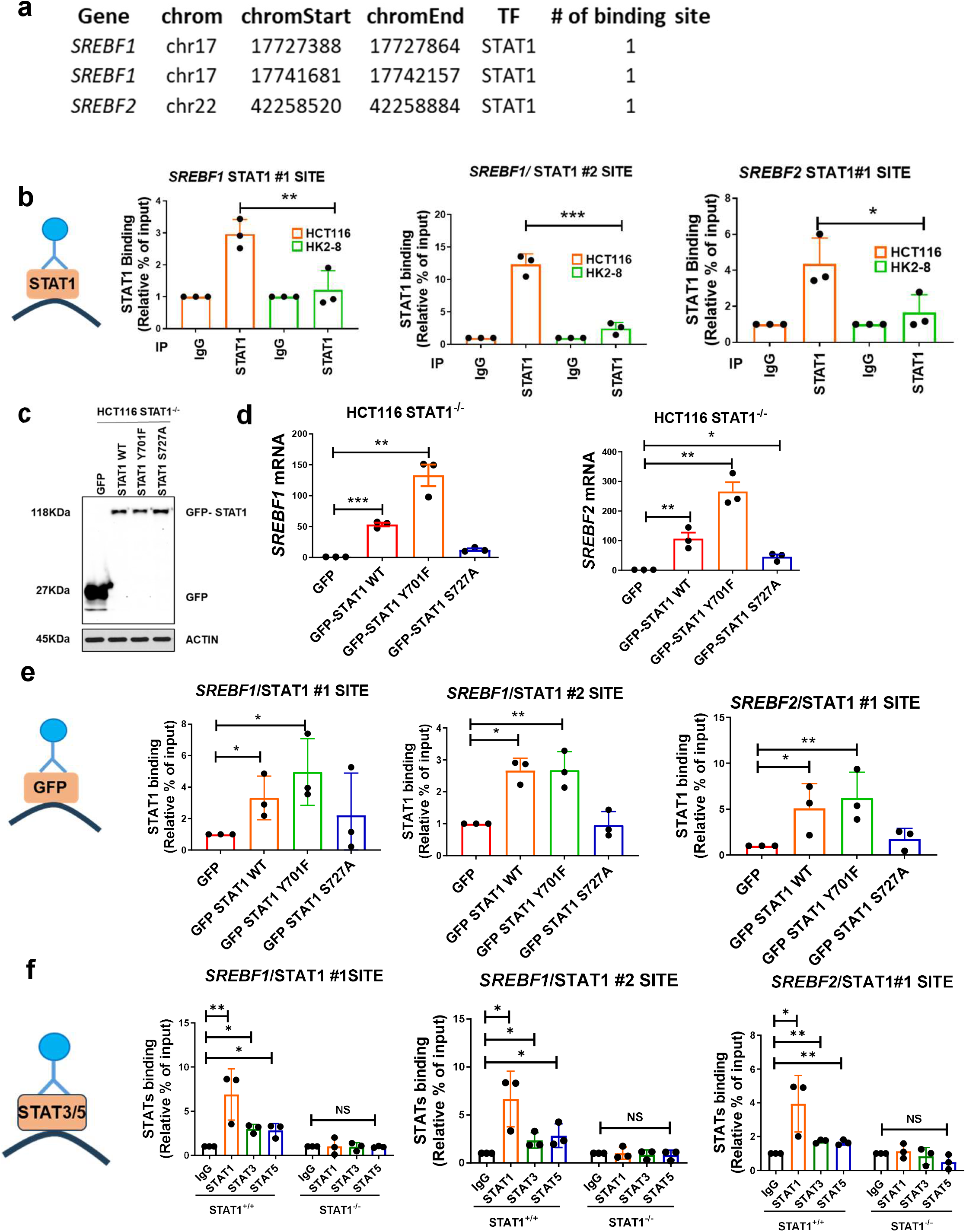
STAT1 stimulates the transcription of SREBF genes in mutant KRAS cells. (**a**) ChIP-seq data from ENCODE (UCSC database) indicating the binding of STAT1 to the regulatory regions of *SREBF1* and *2* genes. (**b**) ChIP assays of endogenous STAT1 bound to *SREBF* gene segments containing STAT1 binding sites in HCT116 and HK2-8 cells. IgG, non-specific control antibody. (**c**) Expression of GFP (control) and GFP-tagged STAT1 proteins that are either intact (wild type, WT) or impaired for phosphorylation (Y701F or S727A) in HCT116 STAT1^-/-^ cells. GFP+ cells were sorted by flowcytometry, and extracts were immunoblotted for GFP or ACTIN. (**d**) Detection of *SREBF-1* and *2* mRNAs by qPCR in HCT116 STAT1^-/-^ cells expressing either GFP or GFP-STAT1 forms. (**e**) ChIP assays of GFP-STAT1 for binding to STAT1 sites of *SREBF* genes in reconstituted HCT116 STAT1^-/-^ cells using GFP antibody. (**f**) ChIP assays of STAT3 and STAT5 for binding to STAT1 sites of *SREBF* genes in HCT116 STAT1^+/+^ and STAT1^-/-^ cells. IgG, non-specific control antibody. (**b-f**) Data obtained from 3 biological replicates and represent ±SEM (*\**) *P* < 0.05 (*\*\**) *P* < 0.01 (***) *P* < 0.001 (*t*-test)

To explore this, we reconstituted HCT116 STAT1^-/-^ cells with green fluorescence (GFP)-tagged forms of human STAT1 that were either intact (WT) or impaired for phosphorylation at Y701 (Y701F) or S727 (S727A). GFP positive cells were sorted and tested for GFP-STAT1 expression via immunoblotting and expression of native *SREBF1* and *SREBF2* genes through qPCR. While GFP-STAT1 proteins were equally expressed in reconstituted STAT1^-/-^ cells (Fig. 3c), *SREBF1* and *2* mRNA levels increased in cells with either GFP-STAT1 WT or GFP-STAT1 Y701F, but not in cells with GFP-STAT1 S727A (Fig. 3d). Subsequent ChIP assays confirmed the binding of GFP-STAT1 WT and GFP-STAT1 Y701F to the regulatory regions of *SREBF1* and *SREBF2* genes in the reconstituted HCT116 STAT1^-/-^ cells (Fig. 3e). In contrast, GFP-STAT1 S727A showed no significant DNA binding compared to cells reconstituted with GFP alone (Fig. 3e). These results suggest that S727 phosphorylation is essential for STAT1’s transcriptional function in upregulating *SREBF* gene expression in mutant KRAS cells.

STAT1 phosphorylation at Y701 enhances its DNA-binding and transcriptional properties in the JAK-STAT pathway, whereas S727 phosphorylation does not affect STAT1’s DNA-binding properties ^37^. Thus, we hypothesized that STAT1 heterodimerization with other STAT family members may account for its ability to bind DNA in the regulatory regions of *SREBF* genes. Through ChIP assays, we observed the binding of STAT3 and STAT5 to *SREBF* gene DNA at sites containing STAT1 binding motifs (Fig. 3f). However, in HCT116 cells lacking STAT1, the binding of STAT3 and STAT5 to DNA was abolished (Fig. 3f). This suggests that STAT3 and STAT5 are involved in facilitating STAT1’s binding to the regulatory regions of *SREBF* genes in HCT116 cells.

### STAT1 promotes the nuclear localization and gene transactivating function of YAP1 in mutant KRAS cells

A notable property of the mevalonate pathway is its ability to enhance the activity of the transcriptional activator YAP1 ^38^, a critical pro-survival factor in mutant KRAS cancers ^16-18^. We hypothesized that STAT1 may, at least partially, rely on YAP1 to exert its pro-survival effects on mutant KRAS CRC cells.

To investigate this, we examined the nuclear trafficking of YAP1 in HCT116 STAT1^+/+^ and STAT1^-/-^ cells treated with either serum or lysophosphatidic acid (LPA), both of which activate YAP1 ^38^. Immunofluorescence analysis demonstrated that both treatments stimulated the nuclear localization of YAP1 in HCT116 cells but not in HK2-8 cells, indicating that mutant KRAS promotes the nuclear trafficking of YAP1 (Fig. 4a). Deletion of STAT1 prevented the stimulation of YAP1 nuclear localization by serum or LPA in HCT116 cells, while in HK2-8 cells, STAT1 inactivation did not produce noticeable effects (Fig. 4a). A similar observation was made when YAP1 nuclear localization was assessed in HCT116 and HK2-8 cells using subcellular fractionation and immunoblotting. Specifically, deletion of STAT1 decreased the nuclear fraction of YAP1 in HCT116 cells but not in HK2-8 cells after serum stimulation (Fig. 4b, c).

**Fig. 4.**
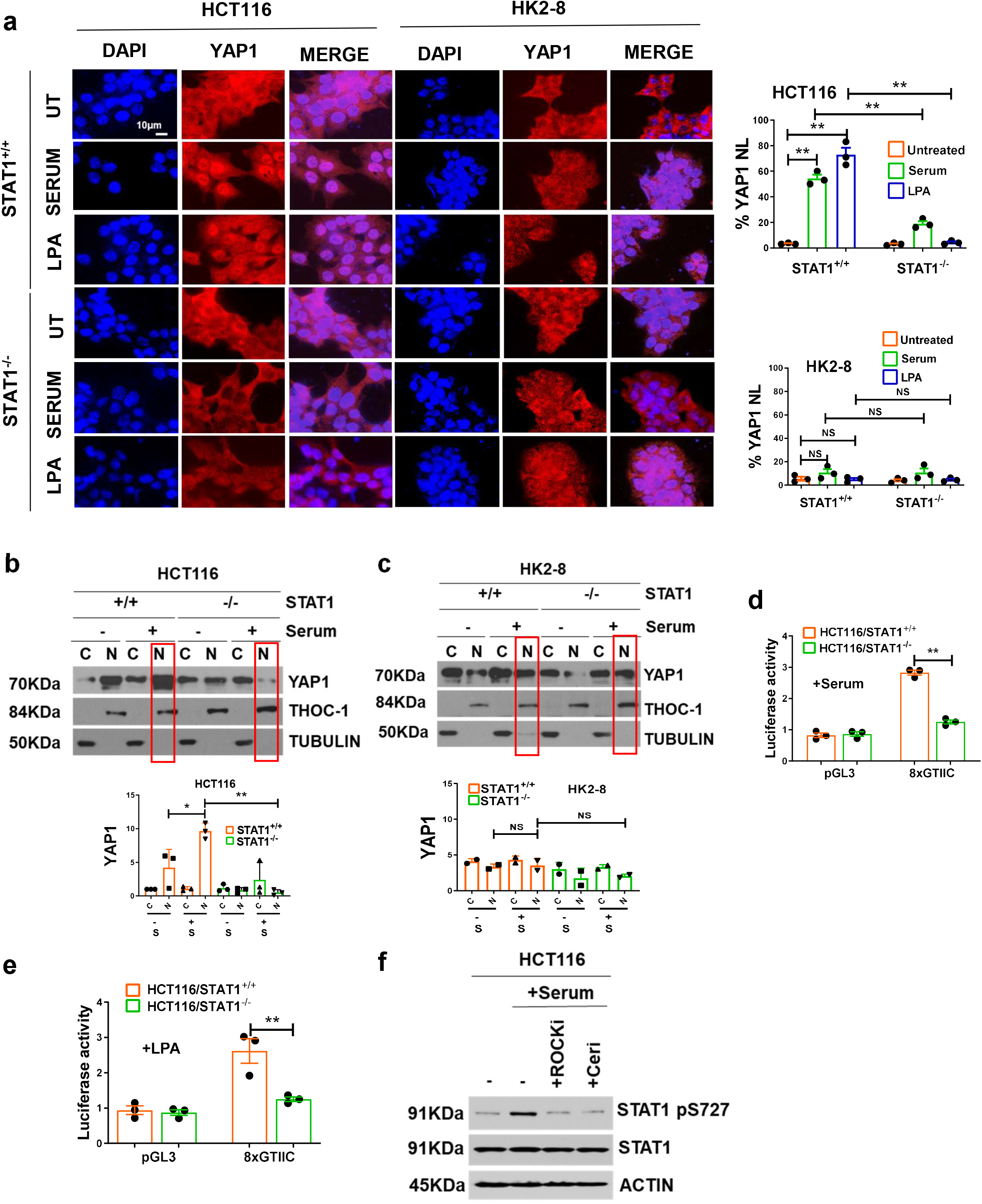
STAT1 stimulates YAP1 in mutant KRAS cells. (**a**) HCT116 and HK2-8 cells replete or deplete of STAT1 were serum-starved for 18 h (untreated; UT) and stimulated with either 10% fetal bovine serum or 25 μM LPA for 1 h. Cells were subjected to IF analyses of YAP1 (red) along with DAPI staining of DNA (blue). Graphs show the quantification of YAP1 nuclear localization in 300 cells. (**b, c**) Cells were subjected to cytoplasmic (C), and nuclear (N) fractionation followed by immunoblotting for the indicated proteins. TUBULIN or THO complex 1 (THOC1) was used as cytoplasmic or nuclear marker, respectively. (**d, e**) HCT116 STAT1^+/+^ and STAT1^-/-^ cells were transfected with either pGL3-luciferase reporter plasmid (control) or 8xGTIIC plasmid containing the firefly luciferase reporter gene under the control of 8x TEAD binding sites in CTGF minimal promoter. Transfected cells were serum starved for 18h followed by stimulation with either 10% fetal bovine serum or 25 μM LPA for 6 h. A plasmid expressing the renilla luciferase gene was used as internal control. (**a**-**e**) Graphs show the quantifications from 3 biological replicates and represent ±SEM (*\**) *P* < 0.05; (*\*\**) *P* < 0.01 (***) *P* < 0.001 (*t*-test), NS, non-significant. (**f**) HCT116 cells were serum starved for 18h followed by stimulation of 10% fetal bovine serum in the absence or presence of 2.5 mM cerivastatin or 10 μM ROCK kinase inhibitor Y-27632 for 18h.

We then investigated the role of STAT1 in YAP1-mediated gene transactivation. Since YAP1 does not have DNA-binding activity, its transcriptional function relies on interactions with DNA-bound transcription factors such as members of the transcriptional enhanced associate domain (TEAD) family ^39^. To assess this, we conducted transient transfection assays using a luciferase reporter gene under the control of 8 binding sites of TEAD in the minimal promoter of the connective tissue growth factor (CTGF) gene (8xGTIIC), a classical YAP1/TEAD-dependent gene ^40^. We found that YAP1-mediated transactivation of the 8xGTIIC promoter was significantly reduced in STAT1^-/-^ compared to STAT1^+/+^ HCT116 cells following serum or LPA treatment (Fig. 4d, e). Taken together, these results support the interpretation that STAT1 promotes the nuclear trafficking and transactivation activity of YAP1 in mutant KRAS cells.

### Activation of YAP1 by STAT1 requires the upregulation of the mevalonate pathway

We further explored whether STAT1’s upregulation of the mevalonate pathway contributes to YAP1 activation in mutant KRAS cells. To this end, we tested the effects of fluvastatin and cerivastatin, which are inhibitors of the rate-limiting enzyme of the mevalonate pathway, HMGCR (54), on YAP1 nuclear localization in HCT116 cells. Immunofluorescence assays revealed that serum stimulation increased YAP1 nuclear localization in HCT116 cells with native STAT1, an effect that was prevented by pre-treatment with either fluvastatin or cerivastatin (Supplementary Fig. 2a). However, YAP1 nuclear localization was not increased by serum in HCT116 STAT1^-/-^ cells either in the absence or presence of fluvastatin or cerivastatin (Supplementary Fig. 2a). The addition of mevalonate metabolite into the media stimulated the nuclear localization of YAP1 in STAT1^+/+^ cells in the presence of fluvastatin or cerivastatin (Supplementary Fig. 2a). This is because addition of mevalonate can bypass the inhibition of the enzymatic activity of HMGCR by fluvastatin or cerivastatin (Fig. 2c). Addition of mevalonate also restored the nuclear localization of YAP1 in STAT1^-/-^ cells in the absence as well as presence of fluvastatin or cerivastatin (Supplementary Fig. 2a). This data verifies that STAT1 depends on the mevalonate pathway to promote the nuclear trafficking of YAP1 in mutant KRAS cells.

Because sterol biosynthesis has been linked to the activation of YAP1 through increased RHO GTPase signaling ^38^, we investigated the effects of Y-27632, an inhibitor of RHO-associated kinases ^41^, on the nuclear localization of YAP1. We observed that while serum stimulation of HCT116 cells increased the nuclear presence of YAP1, treatment with Y-27632 inhibited this process (Supplementary Fig. 2b). Additionally, we found that treating HCT116 cells with either cerivastatin or Y-27632 prevented the serum-induced phosphorylation of STAT1 at S727, implicating the mevalonate pathway and RHO GTPase activation in STAT1 phosphorylation (Fig. 4f).

### YAP1 stimulates the expression of SREBPs and lipid biosynthetic genes in mutant KRAS cells

To understand YAP1’s function, we examined the gene expression profiles of HCT116 cells either with an intact (control) or deleted YAP1 (YAP1^-/-^) by the CRISPR approach. Data analysis revealed that YAP1 promotes the expression of lipid biosynthetic genes (Fig. 5a). Gene ontology and KEGG pathway analyses confirmed that lipid biosynthetic processes, like sterol and fatty acid metabolism, are upregulated by YAP1 in mutant KRAS cells (Fig. 5b). Consistent with YAP1’s role in lipid biosynthesis, the downregulation of YAP1 by siRNAs resulted in a decrease in the mRNA expression of *SREBF1* and SREBF*2*, as well as SREBP-dependent genes such as *ACAT1, HMGCR*, and *IDI1*, in HCT116 cells but not in HK2-8 cells (Fig. 5c; Supplementary Fig. 3).

**Fig. 5.**
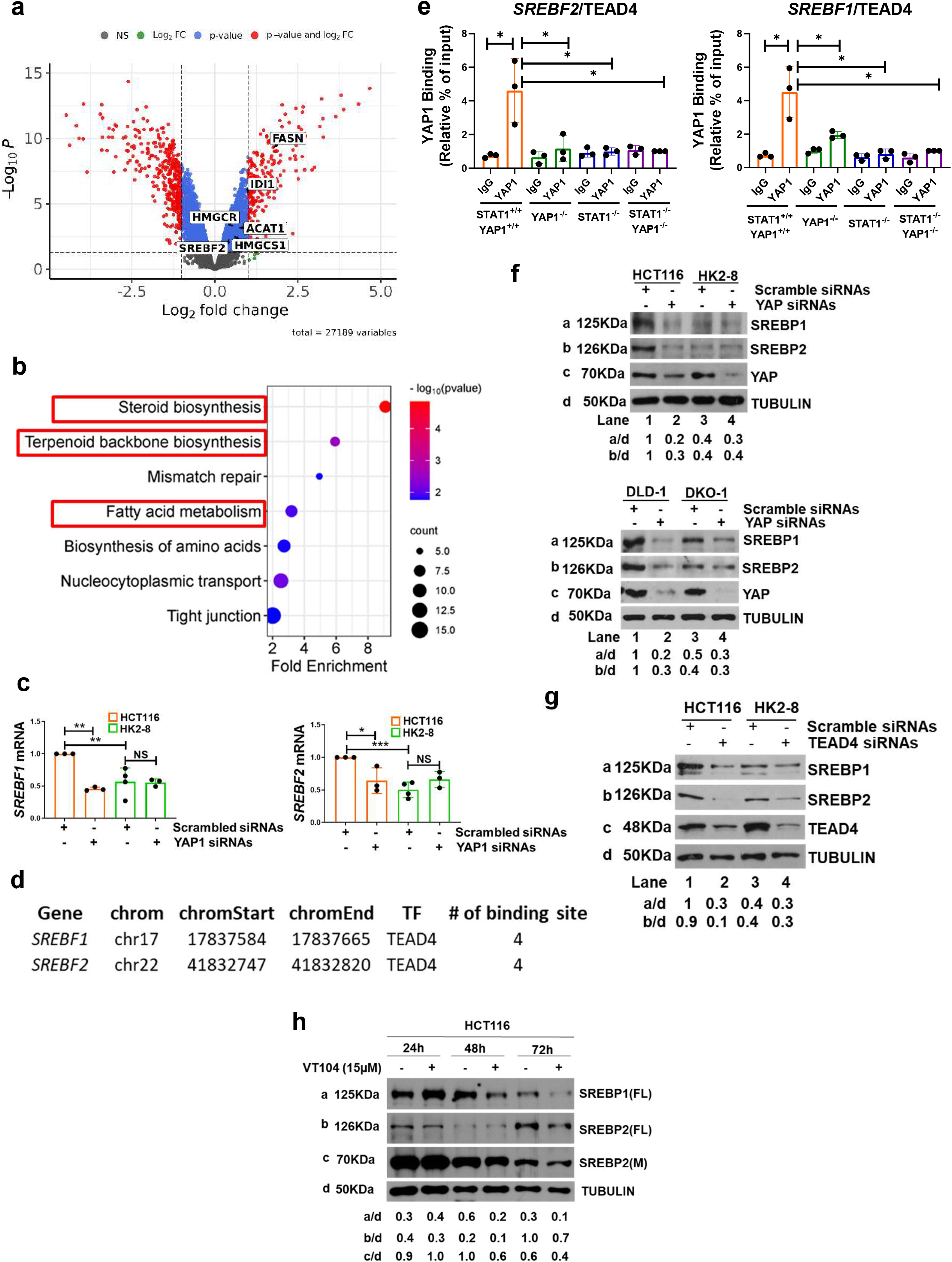
YAP1/TEAD4 upregulates SREBFs and sterol biosynthetic genes in mutant KRAS cells. (**a**) Volcano diagram showing the number of genes differential expressed in YAP1-replete (WT; control) compared to YAP1^-/-^ HCT116 cells. Essential genes of sterol and lipid biosynthetic pathways are indicated. (**b**) Top KEGG pathways under the control of YAP1 in HCT116 cells. (**c**) Graphs assess the expression of *SREBF1* and *2* mRNAs by qPCR in cells treated with scrambled or YAP1 siRNAs. Gene expression was normalized to ACTIN and TUBULIN mRNAs used as internal controls. Data were obtained from 3 independent experiments performed in triplicates and represent ±SEM (*\**) *P* < 0.05; (\**\**) *P* < 0.01;(***) *P* < 0.001 (*t*-test). (**d**) ChIP-seq data from ENCODE indicating the binding of TEAD4 to transcriptional regulatory regions of *SREBF* genes. (**e**) ChIP assays of YAP1 for binding to TEAD4 sites of *SREBF* genes in HCT116 cells, both in the presence and absence of STAT1 and/or YAP1.IgG, non-specific control antibody. (**f**) Immunoblotting of SREBP1 and 2 in isogenic pair colon cancer cells prior to and after YAP1 downregulation by siRNAs. (**g**) Immunoblotting of SREBP1, 2 and TEAD4 in isogenic pairs of colon cancer cells treated with scrambled or TEAD4 siRNAs. (**h**) Immunoblotting of SREBP1 and 2 proteins in HCT116 cells treated with TEAD inhibitor 15 μM VT104 for the indicated time points. (**f, g, h**) Quantification of proteins normalized to TUBULIN for each blot is indicated.

To investigate a possible direct effect of YAP1 on the expression of sterol biosynthetic genes, we analyzed the ChIP-seq data from the ENCODE database. We identified TEAD4 binding sites, a transcriptional partner of YAP1, in the regulatory regions of *SREBF1* and SREBF*2* genes (Fig. 5d). Using ChIP assays, we confirmed YAP1 binding to the regulatory regions of SREBF1 and SREBF2 that contain TEAD4 binding sites (Fig. 5e). This effect was absent in HCT116 cells where YAP1, STAT1, or both proteins were deleted using the CRISPR approach (Fig. 5e). The absence of YAP1 binding in STAT1-deleted cells further supports the role of STAT1 in enhancing YAP1 function in HCT116 cells.

Downregulation of YAP1 by siRNAs resulted in a decrease in SREBP1 and SREBP2 protein levels in HCT116 cells, but not in HK2-8 cells (Fig. 5f). Similarly, siRNA-mediated downregulation of TEAD4 reduced the expression of SREBP1 and SREBP2 proteins in both HCT116 and DLD-1 cells, but not in the isogenic HK2-8 and DKO-1 cells, respectively (Fig. 5g). Additionally, treatment of HCT116 cells with VT104, a potent and specific TEAD inhibitor^42^, led to a reduction in SREBP1 and SREBP2 protein levels in HCT116 cells (Fig. 5h).

These findings strongly support the implication of YAP1-TEAD4 in the upregulation of SREBPs and downstream sterol biosynthetic genes in mutant KRAS cells.

### STAT1 and YAP1 act together via the mevalonate pathway to promote the survival and growth of mutant KRAS cells

Further analysis of the gene expression profiles of proficient and deficient HCT116 cells for STAT1 and/or YAP1 revealed that 186 genes were upregulated by both STAT1 and YAP1 (Fig. 6a). These genes included those with roles in sterol biosynthesis and fatty acid metabolism (Fig. 6d-g), consistent with the ability of STAT1 and YAP1 to stimulate the expression of SREBP1 and 2 at the transcriptional level. Taken together, the data reveal that STAT1 and YAP1 form a feedforward autoregulatory loop that promotes the expression of SREBPs and lipid biosynthetic genes in mutant KRAS cells (Fig. 6e).

**Fig. 6.**
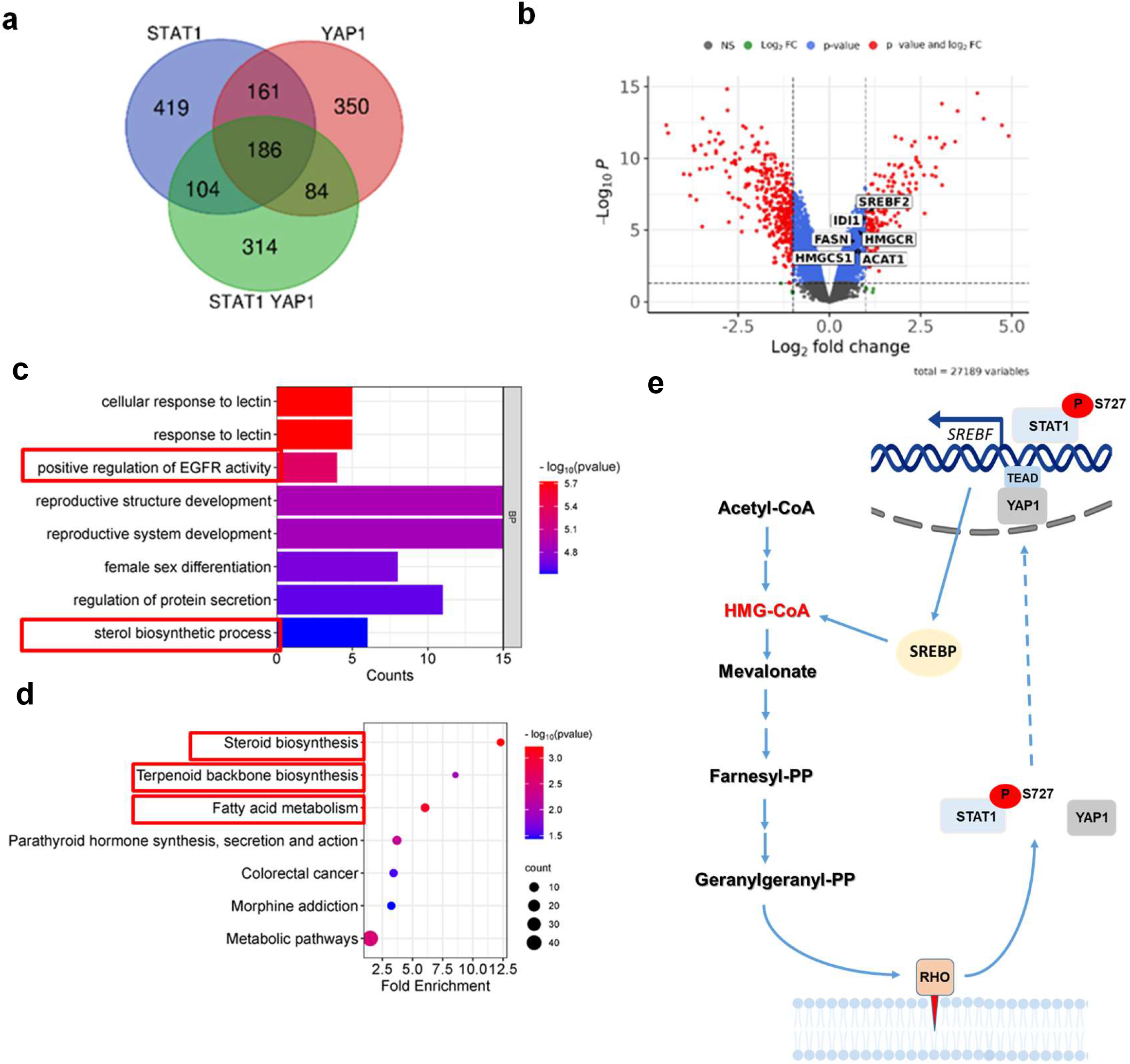
STAT1 and YAP1 act jointly to upregulate SREBPs and lipid biosynthetic pathways. (**a, b**) Venn diagram (a) and Volcano diagram (b) of genes that are commonly upregulated by STAT1 and YAP1 in HCT116 cells. Essential genes of sterol and lipid biosynthetic pathways are indicated in the Volcano diagram. (**c**) Bar plot of biological processes (BP) from gene ontology (GO) significantly enriched in the common set of genes under the control of both YAP1 and STAT1 identified by gene expression profile analysis. (**d**) KEGG pathways under the control of STAT1 and YAP1 in HCT116 cells. (**e**) This schematic illustrates the cooperative role of STAT1 and YAP1 in upregulating SREBPs and the mevalonate pathway. The STAT1-YAP1 axis operates as a feedforward autoregulatory loop that maintains sterol biosynthesis, specifically in mutant KRAS colon cancer cells. The mevalonate pathway activates RHO GTPases, which in turn enhance the phosphorylation of STAT1 at S727 and the nuclear localization and activation of YAP1. While YAP1 functions downstream of STAT1, it also participates in a positive feedback loop that drives the transcription of *SREBF* genes through its interaction with TEAD4.

We next investigated whether the STAT1-YAP1 arm plays a role in the survival and growth of the mutant KRAS cells. Treatment with YAP1 siRNAs notably reduced the ability of HCT116 and DLD-1 cells to form colonies, while it did not affect colony formation in HK2-8 and DKO-1 cells, respectively (Supplementary Fig. 4a). Concurrent treatment with YAP1 and STAT1 siRNAs reduced the survival of HCT116 and DLD-1 cells to a similar degree as individual treatments with either YAP1 or STAT1 siRNAs (Supplementary Fig. 4a). Conversely, treatment with YAP1 and/or STAT1 siRNAs, alone or in combination, did not impact the survival of HK2-8 and DKO-1 cells compared to treatment with control scrambled siRNAs (Supplementary Fig. 4a). Immunoblot analysis confirmed the effectiveness of siRNAs in reducing STAT1 and/or YAP1 levels in the isogenic colon cancer cells (Supplementary Fig. 4b). These findings suggest that STAT1 and YAP1 collaborate to support the survival of mutant KRAS tumor cells.

We further investigated the tumorigenic potential of HCT116 STAT1^-/-^, YAP1^-/-^, and STAT1^-/-^ YAP1^-/-^ cells in immunodeficient mice (Fig. 7a). The deletion of either STAT1 or YAP1 by CRISPR led to a similar decrease in the growth rate of HCT116 tumors in nude mice, and this growth inhibition was not further diminished by the combined deletion of STAT1 and YAP1 (Fig. 7a). This data provides further support for a collaborative function of STAT1 and YAP1 in the stimulation of the growth of mutant KRAS tumors.

**Fig. 7.**
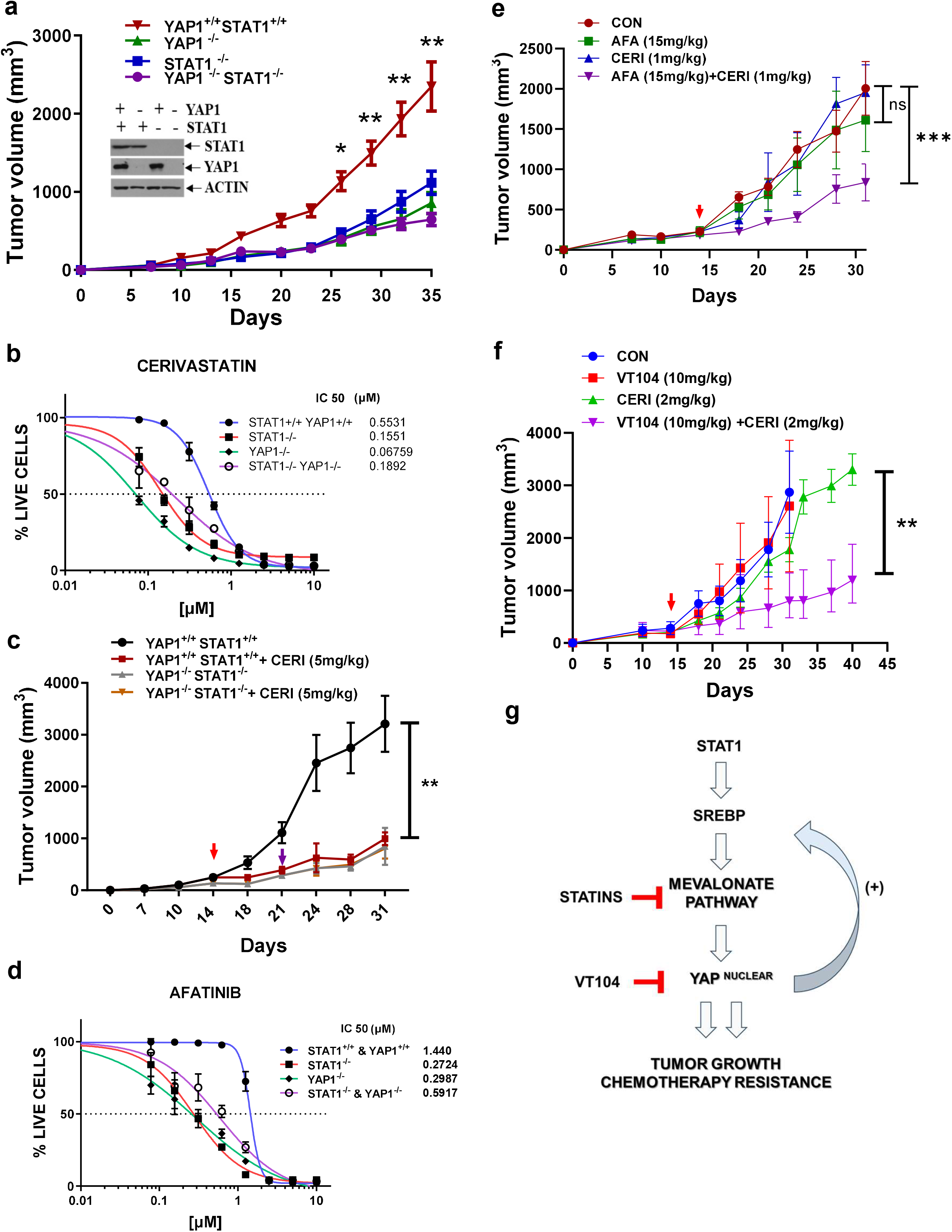
The STAT1-YAP1 axis, through the mevalonate pathway, contributes to the growth of mutant KRAS cells and their resistance to anti-tumor drugs. (**a**) HCT116 cells, both replete and deplete for STAT1 and/or YAP1, were transplanted subcutaneously into female nu/nu mice (n=5). Tumor volume (mm^3^) was measured over the indicated time. Data represent ±SEM (*\**) *P* < 0.05; (**) *P* < 0.01 (*t*-test). (**b**) IC_50_ assays were performed on HCT116 cells that were either replete or deplete for STAT1, YAP1, or both, following treatments with cerivastatin. (**c**) HCT116 cells, either replete or deplete for both STAT1 and YAP1, were similarly transplanted subcutaneously into female nu/nu mice (n=5 per group). Once the tumors, either STAT1/YAP1 replete (red arrow) or STAT1/YAP1 deplete (blue arrow), reached approximately 200 mm^3^, the mice were treated via oral gavage with either vehicle control or cerivastatin (CERI). Tumor volume (mm^3^) was monitored for the indicated period. Data represent ±SEM (**) *P* < 0.01 (*t*-test). (**d**) IC_50_ assays of HCT116 cells that were either replete or deplete for STAT1, YAP1, or both, following treatments to EGFR inhibition with afatinib. (**e**) HCT116 cells were transplanted subcutaneously into nu/nu mice (n=5 per group). Once the tumors reached approximately 200 mm^3^ (red arrow), the mice were treated with either vehicle control, oral gavage of cerivastatin, intraperitoneal administration of afatinib, or a combination of both afatinib and cerivastatin for the indicated times. Tumor volume (mm^3^) was monitored throughout the study. Data represent ±SEM (***) *P* < 0.001 (*t*-test). (**f**) HCT116 cells were transplanted subcutaneously into nu/nu mice (n=5 per group). When the tumors reached approximately 200 mm^3^ (red arrow), the mice were treated with either vehicle control, oral gavage of cerivastatin, oral gavage of VT104, or a combination of both cerivastatin and VT104 for the indicated times. Tumor volume (mm^3^) was monitored throughout the study. Data represent ±SEM (**) *P* < 0.01 (*t*-test). (**g**) This model highlights the biological impact of the STAT1-YAP1 axis in promoting the mevalonate pathway, which drives the growth and increases chemotherapy resistance of CRC with mutant KRAS. Pharmacological inhibition of both the mevalonate pathway and YAP1-TEAD shows potent anti-tumor effects in xenograft mouse models of mutant KRAS CRC.

We addressed the implication of STAT1 and YAP1 in the response of HCT116 cells to mevalonate pathway inhibitors. Through colony formation and IC_50_ assays, we observed that HCT116 cells exhibited increased sensitivity compared to HK2-8 cells when the mevalonate pathway was inhibited by cerivastatin or fluvastatin (Supplementary Fig. 5a,b). This finding suggests that mutant KRAS sensitizes colon cancer cells to mevalonate pathway inhibition. Since neither STAT1 nor YAP1 contributes to the upregulation of the mevalonate pathway or survival of HK2-8 cells, we further investigated their role in determining HCT116 cell responses to treatment with the mevalonate pathway inhibitors cerivastatin and fluvastatin. Colony formation and IC_50_ experiments demonstrated that HCT116 cells with single as well as combined deletion of STAT1 and YAP1 were more vulnerable to mevalonate pathway inhibition than control replete cells (Fig. 7b, Supplementary Fig. 5c,d). These results indicate that the mevalonate pathway plays a major role in the pro-survival effects of STAT1 and YAP1 in mutant KRAS cells.

We assessed the significance of the mevalonate pathway in the tumorigenic effects of STAT1 and YAP1 arm in immune deficient mice. We found that treatment with cerivastatin substantially decreased the growth of control HCT116 tumors proficient for STAT1 and YAP1 (i.e., STAT1^+/+^ YAP1^+/+^ tumors) compared to vehicle treated tumors, and that this growth inhibition was at comparable level to the growth of STAT1^-/-^ YAP1^-/-^ tumors (Fig. 7c). Furthermore, cerivastatin treatment did not lead to additional growth reduction in STAT1^-/-^ YAP1^-/-^ tumors (Fig. 7c), indicating that the mevalonate pathway plays a significant, if not exclusive, role in supporting the tumorigenic function of STAT1 and YAP1 in HCT116 cells harboring mutant KRAS.

### STAT1 and YAP1-mediated resistance to mevalonate pathway inhibition is linked to the efficacy of EGFR and YAP1-TEAD anti-tumor treatments

*KRAS* mutations in CRC play a crucial role in to the development of resistance to epidermal growth factor receptor (EGFR) inhibition via partially understood mechanisms ^43^. Analysis of the STAT1 and YAP1-dependent pathways in HCT116 cells indicated a positive effect of the axis on the regulation of EGFR activity (Fig. 6c). We investigated whether the pro-survival effects of STAT1 and YAP1 influence the response of mutant KRAS cells to EGFR inhibition and whether such an effect could be mediated, at least in part, by the upregulation of the mevalonate pathway.

Examination of the responses of HCT116 cells to treatment with the EGFR inhibitor afatinib revealed that single as well as combined loss of STAT1 and YAP1 renders HCT116 cells more susceptible to treatment than control replete cells (Fig. 7d). Further analysis of the HCT116 cell responses to combined treatments with afatinib with either cerivastatin or fluvastatin showed that mevalonate pathway inhibition synergized with EGFR inhibition to suppress HCT116 cell proliferation (Supplementary Fig. 6a). These anti-proliferative effects of the drug treatments were more pronounced in HCT116 cells with single or combined deletions of STAT1 and YAP1 compared to control HCT116 cells (Supplementary Fig. 6a).

To assess the significance of mevalonate pathway upregulation in treating mutant KRAS tumor cells with EGFR inhibitors, we evaluated HCT116 cell growth in immunodeficient mice before and after treatment with afatinib and cerivastatin. While single treatments with either afatinib or cerivastatin at low concentrations did not yield significant anti-tumor effects, combined treatments significantly suppressed tumor growth rates (Fig. 7e). These results implicate the STAT1-YAP1 axis in the development of resistance to EGFR inhibition via the upregulation of the mevalonate pathway.

Upon further evaluation of HCT116 cells treated with the pan-KRAS inhibitor BI-2865 ^44^, we observed that the cells remained resistant, even in the absence of STAT1 and/or YAP1 (Supplementary Fig. 6b). Additionally, single, or combined treatments of HCT116 cells with inhibitors of the mevalonate pathway, afatinib, BI-2865, or the YAP1-TEAD inhibitor VT104 ^45^ revealed that the most potent antiproliferative effects were achieved with the combined inhibition of the mevalonate pathway and YAP1-TEAD (Supplementary Fig. 6c). To further explore these findings, we tested these treatments in transplantation assays using HCT116 cells in immunodeficient mice. While single treatments with VT104 or low doses of cerivastatin did not significantly inhibit tumor growth, the combination of both treatments resulted in a strong anti-tumor effect (Fig. 7f). Collectively, these results indicate an alternative strategy to EGFR inhibition for impairing HCT116 tumor growth, specifically by targeting the mevalonate pathway and YAP1’s transcriptional function. The findings suggest that inhibition of the STAT1-YAP1 axis with VT104, when combined with mevalonate pathway inhibitors, may effectively suppress mutant KRAS tumor growth (Fig. 7g)

## DISCUSSION

CRC is a heterogenous disease characterized by multiple gene alterations, activation of distinct pathways and metabolic remodeling involved in its pathogenesis ^46^. Large-scale transcriptomic analysis has revealed the molecular heterogeneity of CRC and led to the classification of tumors into four consensus molecular subtypes (CMS). Among these, the CMS3 subtype, representing 32% of patients, is associated with a “metabolic” phenotype enriched in *KRAS* mutations ^47^. *KRAS* mutations in CRC exhibit differential metabolic remodeling, which can impact disease outcomes and responses to anti-cancer treatments. In mouse models of CRC, metabolic profiling accurately stratified tissues based on various combinations of genetic alterations in *APC, KRAS*, and *PTEN* indicating that metabolic profiling can be utilized for patient stratification and the implementation of personalized therapies ^48^.

Previous studies have shown that expression of KRASG13D in colorectal tumor cells alters their metabolic profile compared to WT KRAS, leading to increased cholesterol biosynthesis and suppression of the interferon response ^49^. Consistent with these findings, our research indicates that STAT1 plays a crucial role in promoting the sterol and lipid biosynthetic pathways in HCT116 and DLD1 cells harboring KRASG13D, but not in isogenic tumor cell lines with WT KRAS. We observed that STAT1 stimulates the expression of SREBPs at the transcriptional level leading to the upregulation of *de novo* sterol and lipid biosynthetic genes. In non-tumorigenic mouse models, STAT1 has been found to inhibit energy expenditure in the liver and reduce mitochondrial biogenesis by suppressing the transcription factor PGC1 ^50^. Conversely, PGC1 coactivates SREBPs, enhancing lipogenic gene expression in the mouse liver ^51^. These observations suggest that the upregulation of SREBPs and lipid biosynthetic pathways in our study is not an inherent function of STAT1, but rather a consequence of mutant KRAS expression.

The transcriptional effect of STAT1 on the increased expression of SREBPs is independent of Y701 phosphorylation but relies on its phosphorylation at S727. While Y701 phosphorylation stimulates the DNA-binding and transcriptional properties of STAT1 in the JAK-STAT pathway, unphosphorylated STAT1 at Y701 can also stimulate gene transcription by binding DNA with low affinity or interacting with other DNA-binding proteins ^37^. Our ChIP assay findings indicate that STAT3 and STAT5 contribute to STAT1’s ability to bind DNA in the regulatory regions of *SREBF* genes. Conversely, S727 phosphorylation can enhance the transcriptional functions of STAT1, independent of Y701 phosphorylation, in cells exposed to UV, IL1 or TNF ^37^. In Wilm’s tumors, one of the most common pediatric solid cancers, STAT1 S727 phosphorylation stimulates tumor growth and protects tumor cells from apoptosis under conditions of stress ^52^. Our research has demonstrated that STAT1 S727 phosphorylation in KRAS G13D tumor cells is constitutively upregulated via the mevalonate pathway, necessitating the involvement of RHO GTPases. We found that pharmacological inhibition of RHO-associated kinase significantly reduces STAT1 S727 phosphorylation, suggesting the involvement of this kinase or another downstream kinase in this process. While RHO GTPases have been previously shown to mediate the phosphorylation and transcriptional activation of STAT3 and STAT5^53^, their role in stimulating STAT1 S727 phosphorylation in mutant KRAS cells represents a novel finding. This insight could potentially lead to new anti-tumor strategies targeting the phosphorylation of STAT1 for cancers harboring *KRAS* mutations.

Although previous studies have identified a role for RHO GTPases in stimulating the nuclear localization and function of YAP1^38^, our data reveal that this process is specifically controlled by STAT1 in KRAS G13D colon cancer cells. The stimulation of YAP1 function by STAT1 relies on the upregulation of the mevalonate pathway, representing a feedforward loop that sustains the upregulation of sterol biosynthetic genes through the transcriptional activation of *SREBF* genes. In non-tumorigenic models, YAP1 upregulates serum- and glucocorticoid-regulated kinase (SGK) 1 to stimulate mTOR activity, leading to SREBP activation ^28^, Additionally, YAP1 directly interacts with SREBPs to promote lipid biosynthetic gene transcription in hepatocytes of diabetic mice ^29^. However, in KRAS G13D colon cancer cells YAP1 connects with STAT1 to sustain SREBP expression and lipid biosynthesis, highlighting a unique interplay between STAT1, YAP1, and the mevalonate pathway in cancer cell metabolism and growth.

Our study demonstrates that the STAT1-YAP1 axis, through the upregulation of the mevalonate pathway, significantly enhances the growth of mutant KRAS tumors. This aligns with other research indicating a tumorigenic role for SREBPs in mutant *KRAS* cancers and the positive effects of mevalonate inhibition by statins on the prevention and treatment of CRC^54^. This is consistent with other research that highlights the tumorigenic role of SREBPs in mutant KRAS-driven cancers ^55,56^and the beneficial impact of mevalonate pathway inhibition by statins in CRC prevention^54^. Additionally, STAT1 and YAP1 jointly confer resistance to mevalonate pathway inhibitors in mutant KRAS cells. Although STAT1 has been previously recognized for its role in protecting tumor cells from chemotherapeutic drugs ^57,58^, this is the first report highlighting its involvement in mediating tumor survival against metabolic therapies targeting a mutant KRAS cancer. On the other hand, multiple studies have supported a pro-survival role for YAP1 in mutant KRAS cancers ^16-19^. Inhibiting the transcriptional activity of YAP1 with TEAD inhibitors significantly disrupts mutant KRAS effector pathways and sensitizes tumor cells to drugs targeting mutant KRAS ^59-61^. We observed that HCT116 cells are resistant to pharmacological inhibition of mutant KRAS but highly sensitive to mevalonate pathway inhibitors especially when combined with the pharmacological inhibition of YAP1-TEAD. Thus, *KRAS* mutations may shift the dependency of tumors to other pathways for their growth like the mevalonate pathway. Consistent with this interpretation, previous work demonstrated that several secondary mutations can contribute to a switch in oncogene addiction in mutant KRAS cancers, an effect that is associated with the activation of distinct effector pathways and metabolic alterations that can determine specificity to anti-tumor treatments ^62^.

In CRC, mutant KRAS is recognized as a key factor influencing response to chemotherapeutic therapies ^43^. For instance, resistance to anti-EGFR therapy is closely associated with *KRAS* mutations through mechanisms that are not fully understood ^43^. Activation of YAP1 by the mevalonate pathway has been associated with resistance to EGFR inhibition in colorectal cancer ^63^. Expression of KRASG13D in CRC cells alters EGFR networks, reshaping the metabolic and transcriptional landscapes of tumor cells, which has significant implications for therapy ^49,64^. Our findings indicate that the STAT1-YAP1 axis plays a role in the development of resistance to the EGFR inhibitor afatinib by upregulating the mevalonate pathway. Notably, this resistance can be overcome by inhibiting the mevalonate pathway in xenograft assays in mice.

The cooperation between STAT1 and YAP1 in promoting lipid biosynthesis may have significant implications for the establishment of anti-tumor immunity in mutant KRAS cancers. STAT1 is a key player in the IFN response triggered by the activation of the stimulator of interferon genes (STING) pathway ^65^. In contrast, YAP1 has antagonistic effects on STING pathway activation and anti-tumor immunity ^66^. However, research has shown that cholesterol biosynthesis can undermine STING signaling in virus-infected cells and tumor cells ^67-69^. Cholesterol also promotes the retention of STING in the endoplasmic reticulum, a process that can be reversed by cholesterol depletion, thereby stimulating anti-tumor immunity when treated with immune checkpoint inhibitors ^70^. The ability of STAT1 to collaborate with YAP1 to upregulate the mevalonate pathway may reveal a new mechanism used by mutant KRAS to compromise the IFN response and impair anti-tumor immunity. This potential function of STAT1 would differ from its established role in activating the IFN response and could provide a crucial link between its tumor-intrinsic and immune regulatory functions in the treatment of mutant KRAS cancers.

## MATERIALS AND METHODS

### Cell culture and treatments

HCT116, HK2-8, DLD-1, and DKO-1 cells were cultured in Dulbecco’s Modified Eagle Medium (Wisent) supplemented with 10% fetal calf serum (Wisent) and 100 U/mL penicillin-streptomycin (Wisent). Isogenic colon cancer cell lines were authenticated for the presence of the *KRAS G13D* by DNA sequencing. Depletion of STAT1 and YAP1 using clustered regularly interspaced palindromic repeats (CRISPR)/CAS9 was conducted as previously described ^9,71^; the guide RNA sequences are provided in Supplementary Table 1. Control cells were those expressing CRISPR/Cas9 vectors lacking guide RNA sequences. Transient downregulation of STAT1 or YAP1 was achieved using four different siRNAs (Horizon Discovery) with sequences listed in Supplementary Table 1. Fluvastatin, cerivastatin, afatinib and BI-2865 were purchased from MedChemExpress, VT104 was acquired from DC Chemicals, Y-27632 was sourced from Selleckhem and 4’,6-diamidino-2-phenylindole (DAPI) was obtained from Roche. Colony formation assays were conducted with 2×10^3^ cells in 6-well petri dishes subjected to treatments as indicated in the Fig. legends for up to 14 days. Cells were fixed in 3.7% formaldehyde (v/v) and stained with 0.2% crystal violet (w/v). Colonies were scored using an automated cell colony counter (GelCount; Oxford Optronix).

### siRNA, DNA transfections and luciferase reporter assay

Plasmid and siRNA transfections were carried out using Lipofectamine 2000 reagent (Invitrogen) according to the manufacturer’s specifications. The 8xGTIIC-luciferase construct was obtained from Addgene (plasmid #34615)^40^. Plasmids containing the green fluorescence protein (GFP)-tagged forms of wild type STAT1, STAT1 Y701F or STAT1 S727A cDNA were obtained from Addgene (plasmid #12301,12302 and 12304)^52^. *Firefly* luciferase assays were performed with the Dual-Luciferase Reporter Assay System (Promega) using *Renilla* luciferase reporter gene serving as an internal control ^72^.

### Chromatin immunoprecipitation (ChIP) and real-time PCR

ChIP assays were conducted following previously established protocols ^72^. Briefly, cells were cross-linked with formaldehyde at a final concentration of 1% for 10 min followed by addition of glycine in PBS at a final concentration of 125 mM. Cells were rinsed ice cold 1XPBS twice and scraped and collected in 1 ml ice cold PBS. Cells were spun down and washed with cold PBS once, and nuclear-enriched extracts were prepared by 1 ml lysis buffer (5mM PIPES, pH 8.0, 85Mm KCl, 0.5% NP-40) plus protease inhibitor. The lysate was sonicated with a Sonifier (Vibra Cell, Soinc & Material INS) to shear the chromatin (Output 20%, 100% duty cycle, 8×15-sec pulses), and the samples were clarified by centrifugation. Protein–DNA complexes were immunoprecipitated (IP) with anti-STATs or GFP-Trap Agarose overnight at 4°C on a rotator. Anti-STATs IP samples were incubated with protein A-agarose for 1 h at 4°C. Anti-STATs or GFP-Trap Agarose samples were followed by one time low salt wash buffer (0.1% SDS, 1% Triton X-100, 2mM EDTA, 20mM Tris-Cl pH8.1, 150mM NaCl), two time high salt wash buffer (0.1% SDS, 1% Triton X-100, 2mM EDTA, 20mM Tris-Cl pH8.1, 500mM NaCl), one time LiCl buffer (250mM LiCl, 1% NP40, 1mM EDTA, 10mM Tris-CL pH8.1, 1% deoxycholate). The protein-DNA complexes were eluted from the beads by addition of 1% SDS, 0.1 M NaHCO_3_ at 65 °C for 1 hour. The beads were removed by centrifugation, and the supernatants were incubated overnight at 65°C to reverse the cross-linking. The DNA was extracted by the addition of RNase A (50μg/ml, 37°C for 30min) and proteinase K solution (300 µg/mL, 55°C for 2 hours) and subjected to purification with a QIAquick PCR purification kit. The enriched DNA was quantified by real-time PCR using a set of primers that cover the STAT1-binding site in the SREBF1 and SREBF2 genes (Supplementary Table 1). Signals obtained from each immunoprecipitation were expressed as a percent of the total input chromatin. Percent (%) input=2% x 2 ^(C[T]2%Input Sample-c[T]ISample)^ where C[T]=C_T_= threshold cycle of PCR reaction.

Total RNA was isolated using TRIzol (Invitrogen), and 1 µg of RNA was reverse transcribed (RT) using the SuperScript III Reverse Transcriptase kit (Invitrogen) with 100 µM oligo (dT) primer, following the manufacturer’s instructions. Real-time PCR was carried out using the SensiFast SYBR Lo-ROX kit (Bioline) with primers listed in Supplementary Table 1. The PCR assays included primers for *GAPDH* and *ACTIN* mRNAs as internal controls, in accordance with the Minimum Information for Publication of Quantitative Real-Time PCR Experiments (MIQE) guidelines ^73^.

### Immunoblotting

Cells were washed twice with ice-cold PBS, and proteins were extracted using an ice-cold lysis buffer containing 10 mM Tris-HCl (pH 7.5), 50 mM KCl, 2 mM MgCl2, 1% Triton X-100, 3 μg/ml aprotinin, 1 μg/ml pepstatin, 1 μg/ml leupeptin, 1 mM dithiothreitol, 0.1 mM Na3VO4, and 1 mM phenylmethylsulfonyl fluoride. The extracts were incubated on ice for 15 minutes, then centrifuged at 10,000 × g for 15 minutes at 4 °C. Supernatants were stored at −80 °C. Protein concentrations were measured using the Bradford assay (Bio-Rad). To evaluate the expression of various proteins, 50 µg of protein extracts from the same set of samples were loaded in parallel onto two identical sodium dodecyl sulfate (SDS)-polyacrylamide gels. After transferring the proteins to Immobilon-P membranes (Millipore), the blots were cut into smaller sections based on the molecular weights of the target proteins. One section was probed for the phosphorylated protein of interest, while the corresponding section was probed for the total protein. The antibodies used for immunoblotting are listed in Supplementary Table 2. Protein visualization was performed using enhanced chemiluminescence (ECL) according to the manufacturer’s instructions (Amersham Biosciences). Band quantification within the linear range of exposure was carried out using ImageJ 1.51e software (NIH, Maryland, USA).

### Gene expression analysis

Samples were processed using the Affymetrix Clariom S Human HT Expression platform. The data was normalized using the ‘rma’ method in the Bioconductor package ‘oligo’. The ‘rma’ method involves background subtraction, normalization with the RMA algorithm, and summarization using median-polish. The expression values were then transformed to the log2 scale. Annotation for the Affymetrix Clariom S Human HT was obtained using the ‘clariomshumanhttranscriptcluster.db’ database provided by Bioconductor. Differential expression analysis of genes in STAT1 and/or YAP1-deficient cells compared to proficient (control) cells was performed using the Bioconductor package ‘limma’ ^74^. Gene network analysis was conducted using the output of the Gene Ontology (GO) analyses with the Bioconductor package ‘DOSE’ ^75^.

### Subcellular fractionation and immunofluorescence assays

Subcellular fractionation and immunofluorescence were performed as described in the protocol ^38^. Cells were lysed on 10 cm plates using 500 µl of buffer containing 250 mM sucrose, 20 mM HEPES, 10 mM KCl, 1.5 mM MgCl2, 1 mM EDTA, and 1 mM EGTA, and were incubated on ice for 20 minutes. The nuclear pellet was obtained by centrifugation at 720g for 5 minutes, then washed and lysed with standard RIPA lysis buffer. The supernatant, representing the cytoplasmic fraction, was collected, and sample buffer was added. THOC-1 served as the nuclear marker, while tubulin was used as the cytoplasmic marker.

For immunofluorescence staining, cells were fixed with 4% paraformaldehyde for 10 minutes, washed with PBS, permeabilized with 0.5% Triton X-100 for 5 minutes, and blocked with 5% goat serum and 5% BSA for 30 minutes. Antigen detection was carried out by incubating the cells with the primary antibody overnight at 4°C, followed by incubation with goat anti-rabbit Alexa Fluor 546 as the secondary antibody for 30 minutes at room temperature. Nuclei were counterstained with 0.5% DAPI. The primary and secondary antibodies used are listed in Supplementary Table 2.

### Xenograft tumor assays

Tumor transplantation assays were conducted in 8-week-old female athymic nude mice (Charles River Inc.) following the established protocol ^9^. Briefly, cells were suspended in a 1:1 v/v mixture of phosphate-buffered saline (PBS): Matrigel (Corning) and injected subcutaneously into the flanks of nude mice. Mice were inoculated with 5×10^5^ cells in 200 µl PBS: Matrigel mix. Tumor growth was monitored twice per week using digital calipers, and the volume was calculated using the formula: tumor volume [mm^3^] = π/6 × (length [mm]) × (width [mm]) × (height [mm]). Mice were subjected to treatments with afatinib, cerivastatin or VT104 at doses reported to have minimal side effects on body toxicity ^59,76-78^. The animal studies were conducted in accordance with the Institutional Animal Care and Use Committee (IACUC) of McGill University, and all procedures were approved by the Animal Welfare Committee of McGill University (protocol #5754).

### Statistical analysis

Statistical analysis was conducted using three biological replicates. Error bars represent the standard error as indicated, and significance in differences between arrays of data was determined using a two-tailed Student’s t-test.

## ACKNOWLEDGEMENTS

We thank S. Shirasawa (Fukuoka University, Japan) for HCT116 and HK2-8 cells ; M. Sudol (National University of Singapore) for the CRISPR-YAP1_gRNA construct; R. DeBose-Boyd (UT Southwestern Medical Center) for HMGCR (A9) mouse monoclonal antibody. The work was supported by funds from the Cancer Research Society Inc., the Marjorie Sheridan Innovation grant of the Canadian Cancer Society (CCSRI #701631) and Canadian Institutes of Health Research (CIHR; PJT-178173) to AEK. HK is supported by George G. Harris Fellowship in Cancer Research from McGill university and internal scholarship award from *Lady Davis* Institute. JYZ is supported by a studentship from *Fonds de recherche Santé Québec*.

## AUTHORS CONTRIBUTIONS

SW collected, analyzed, interpreted the data and prepared the manuscript; SD performed the bioinformatics analyses of the sequencing data; HK assisted in transfection assays of cells; KKL contributed to analysis of cell responses to drug treatments and JYZ assisted in the analysis of anti-tumor treatments in mice; AEK designed the study, interpreted the data, wrote, and edited the manuscript.

## DATA AVAILABILITY

The RNA seq data generated in this study have been deposited in the Gene Expression Omnibus (GEO) GSE278710

