## Supplementary material for "A feedforward loop between STAT1 and YAP1 stimulates lipid biosynthesis, accelerates tumor growth, and promotes chemotherapy resistance in mutant KRAS colorectal cancer": SUPPL FIGURES TABLES

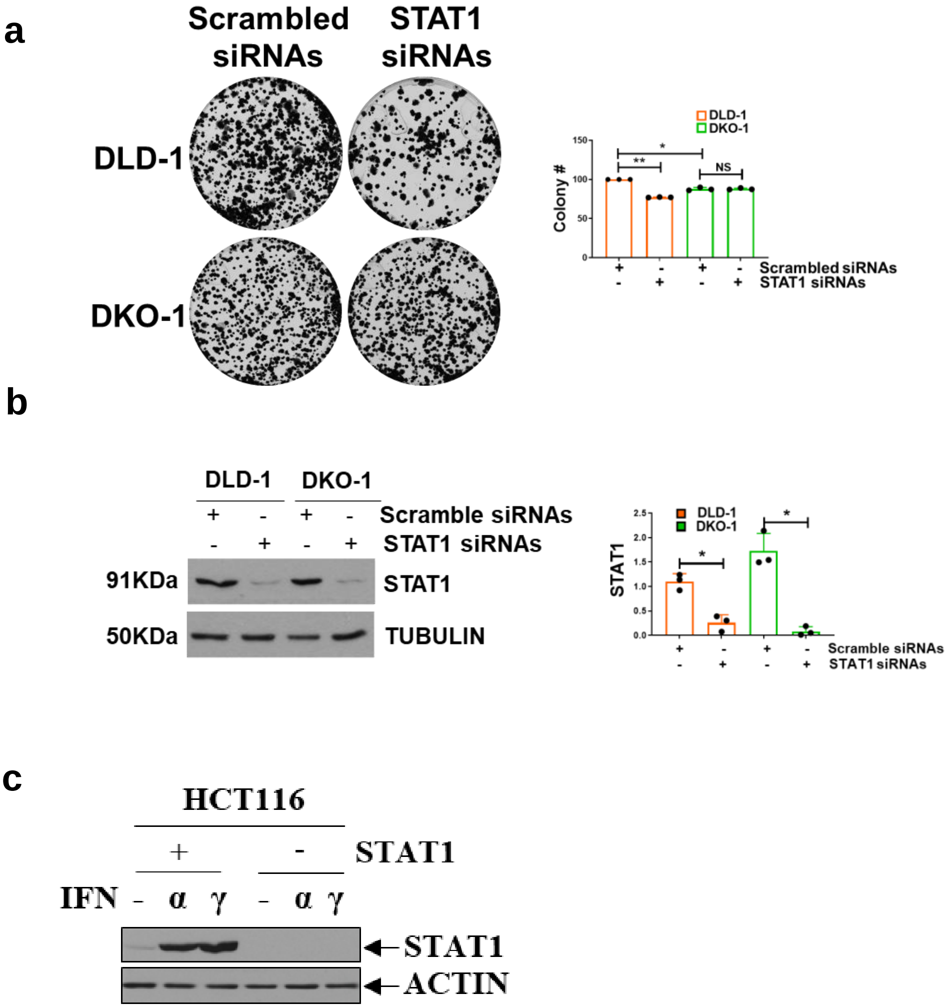

**Supplementary Fig. 1**. ***STAT1 promotes the survival of colon cancer cells with mutant KRAS***. (**a**) Assessment of colony formation efficacy of DLD-1 and DKO-1 cells after treatments with scramble or STAT1 siRNAs. (**b***)* Immunoblotting for STAT1 in DLD-1 and DKO-1 cells treated with . scramble or STAT1 siRNAs. (**a**, **b**) The data were obtained from 3 biological replicates each of which included 3 technical replicates and represent ±SEM (*) $P$ < 0.05 (**) $P$ < 0.01;(*t*-test), NS=non-significant. (**c***)* Immunoblotting for the detection of STAT1 in HCT116 cells either replete or deplete for STAT1 using CRISPR technology. Cells were treated with 500 IU of type I (α) or type II (γ) IFN for 18h.

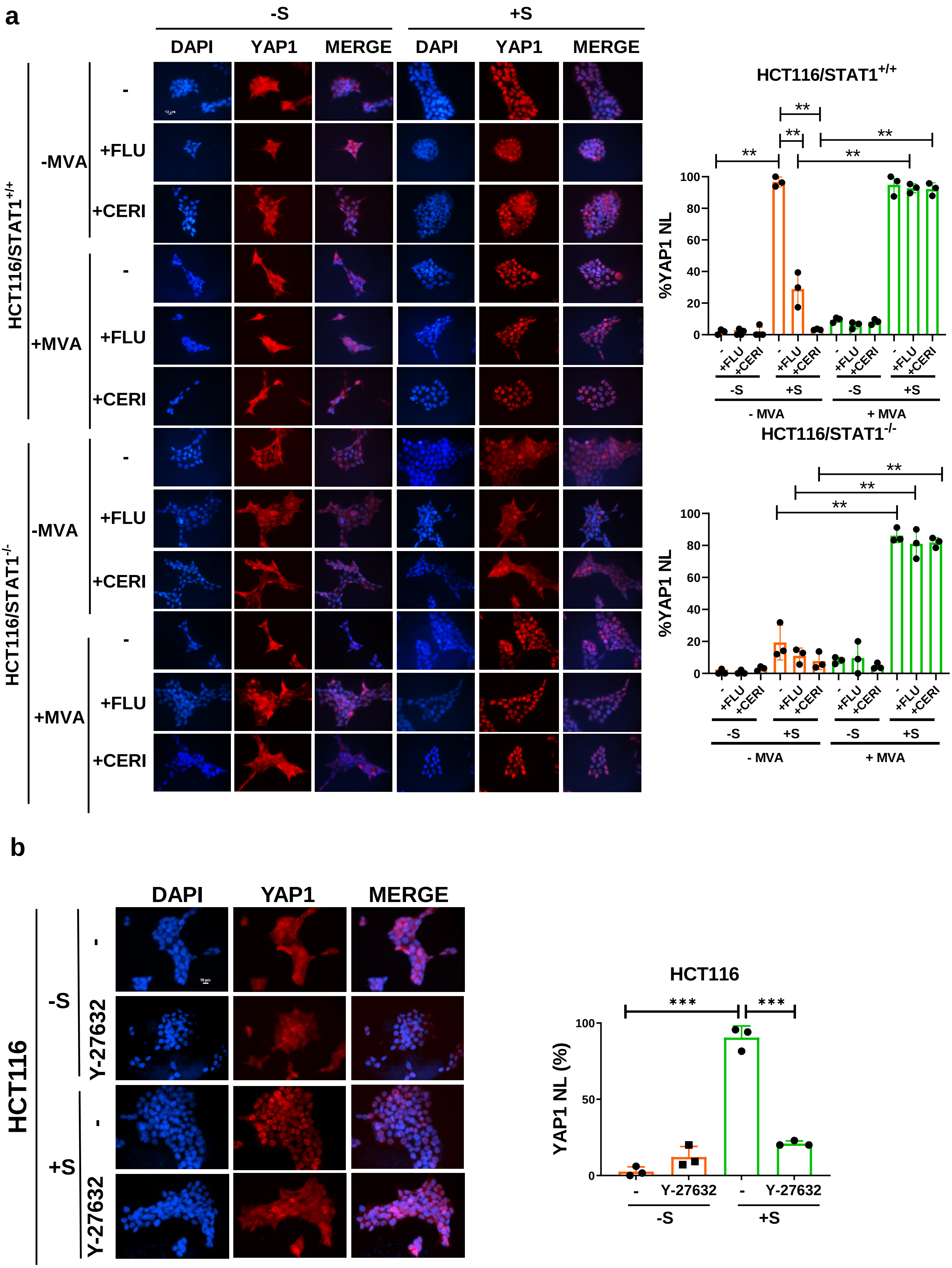
a
-S
+S
DAPI
YAP1
MERGE
DAPI
YAP1
MERGE
HCT116/STAT1+/+
-MVA
+MVA
-
+FLU
+CERI
HCT116/STAT1-/-
HCT116/STAT1+/+
%YAP1 NL
**
- MVA
+ MVA
HCT116/STAT1-/-
b
HCT116
Y-27632
HCT116
YAP1 NL (%)
***

**Suppl Fig. 2**. ***STAT1 promotes YAP1 nuclear localization via the stimulation of the mevalonate pathway.*** (**a**) HCT116 cells replete or deplete of STAT1 were serum-starved for 18 h. Cells were incubated with either 5 µM Fluvastatin or 2.5 µM cerivastatin for 18h followed by stimulation with 10% fetal bovine serum for 1h. A similar set of cells was subjected to the same treatments with the addition of 0.5 mM mevalonate in the media for 18h. Cells were subjected to IF of YAP1 (red) along with DAPI staining of DNA (blue). Symbols are as follows: **-**, untreated cells; **-S**, serum-starved cells ; **+S**, serum-stimulated cells for 1h; **+FLU -S**, serum-starved cells treated with Fluvastatin for 18h; **+FLU +S**, serum-starved cells treated with fluvastatin for 18h and stimulated with serum for 1h; **+CERI -S**, serum-starved cells treated with cerivastatin for 18h; **+CERI +S**, serum-stimulated cells treated with cerivastatin for 18h and stimulated with serum for 1h; **MVA**, cells treated with mevalonate for 18h and maintained in media during serum stimulation. (**b**) HCT116 cells were serum-starved for 18 h. Cells were incubated with 10 µM ROCK kinase inhibitor Y-27632 for 18h followed by stimulation with 10% fetal bovine serum for 1h. (**a**, **b**) The graph shows the quantification of YAP1 unclear localization (NL) in 300 cells from 3 biological replicates and represent ±SEM (**) $P < 0.01$; (***) $P < 0.001$ (*t*-test).

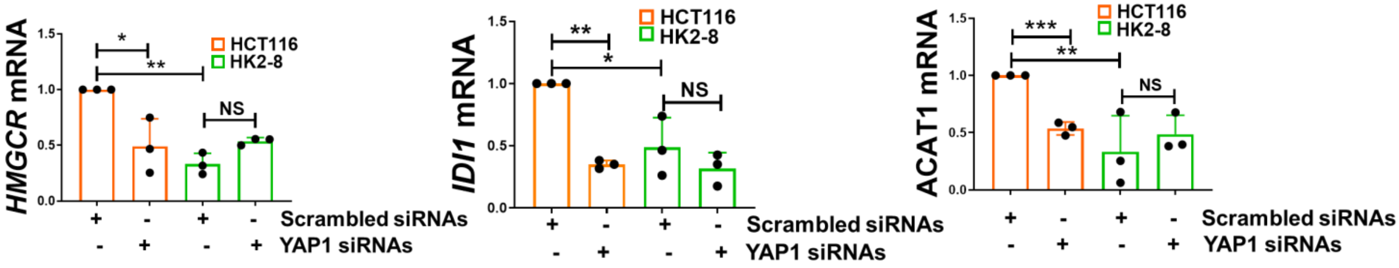

**Suppl Fig. 3**. ***YAP1 mediates the upregulation of mevalonate pathway genes in mutant KRAS cells***. Graphs show the detection of *ACAT1*, *HMGCR* and *IDI1* mRNAs by qPCR in cells treated with scrambled or YAP1 siRNAs. Gene expression was normalized to ACTIN and TUBULIN mRNAs used as internal controls. Data were obtained from 3 independent experiments performed in triplicates and represent ±SEM (*) $P < 0.05$; (**) $P < 0.01$; (***) $P < 0.001$ (*t*-test).

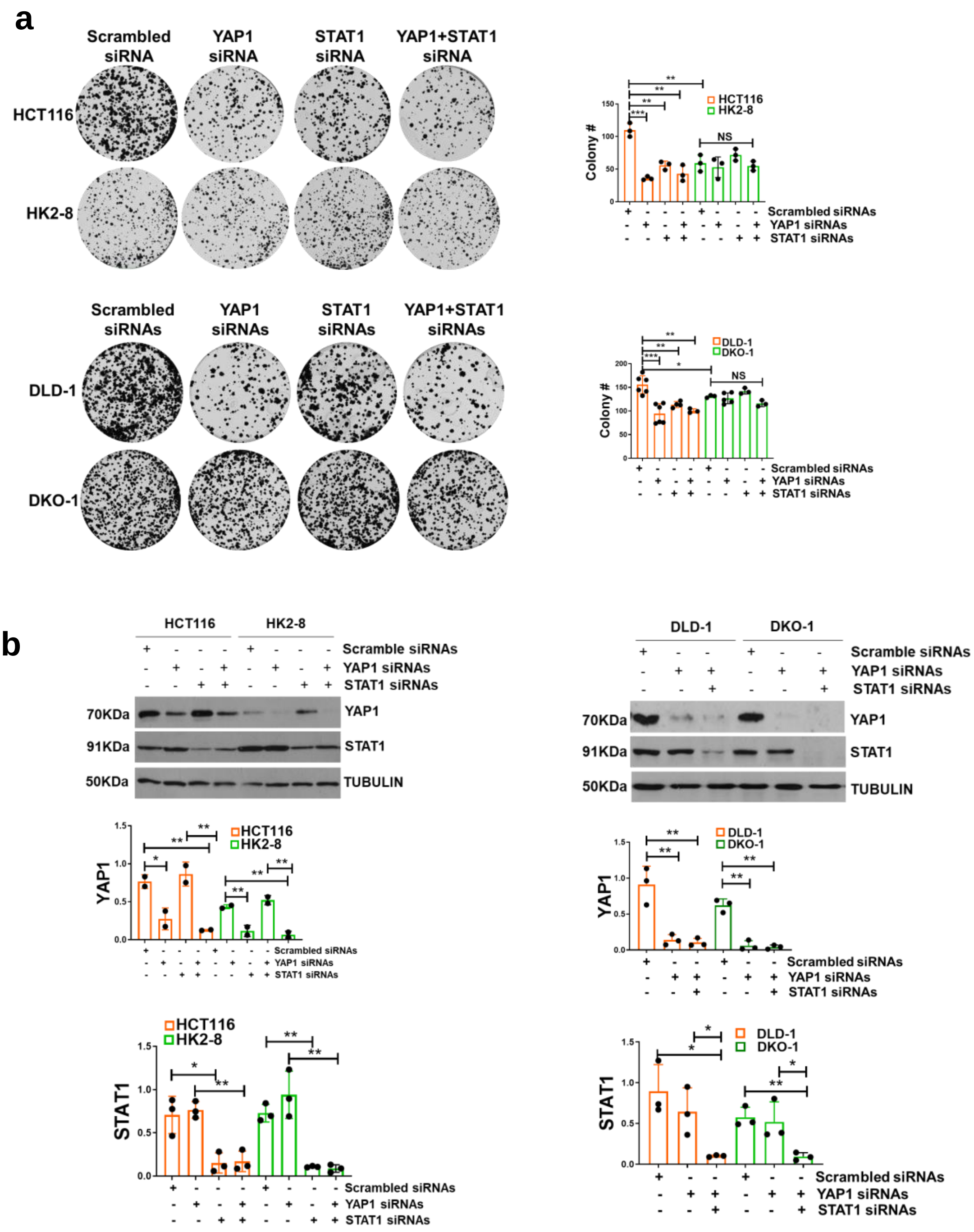

**Suppl Fig. 4.** ***STAT1 and YAP1 cooperate in the stimulation of survival of mutant KRAS cells***. (**a**) HCT116 and HK2-8 cells together with DLD-1 and DKO-1 cells were subjected to colony formation assays under conditions in which expression of YAP1, STAT1, STAT1 and YAP1 was impaired by siRNAs. The graphs represent data obtained from 3 biological replicates each of which included 3 technical replicates and represent ±SEM (*) $P$ < 0.05; (**) $P$ < 0.01; (***) $P$ < 0.001 (*t*-test), NS=non-significant. (**b**) Immunoblotting for STAT1 and YAP1 expression in isogenic colon cancer cells treated with siRNAs. Quantifications show the relative intensity of STAT1 normalized to TUBULIN from 3 biological replicates and represent ±SEM (*) $P$ < 0.05; (**) $P$ < 0.01 (*t*-test).

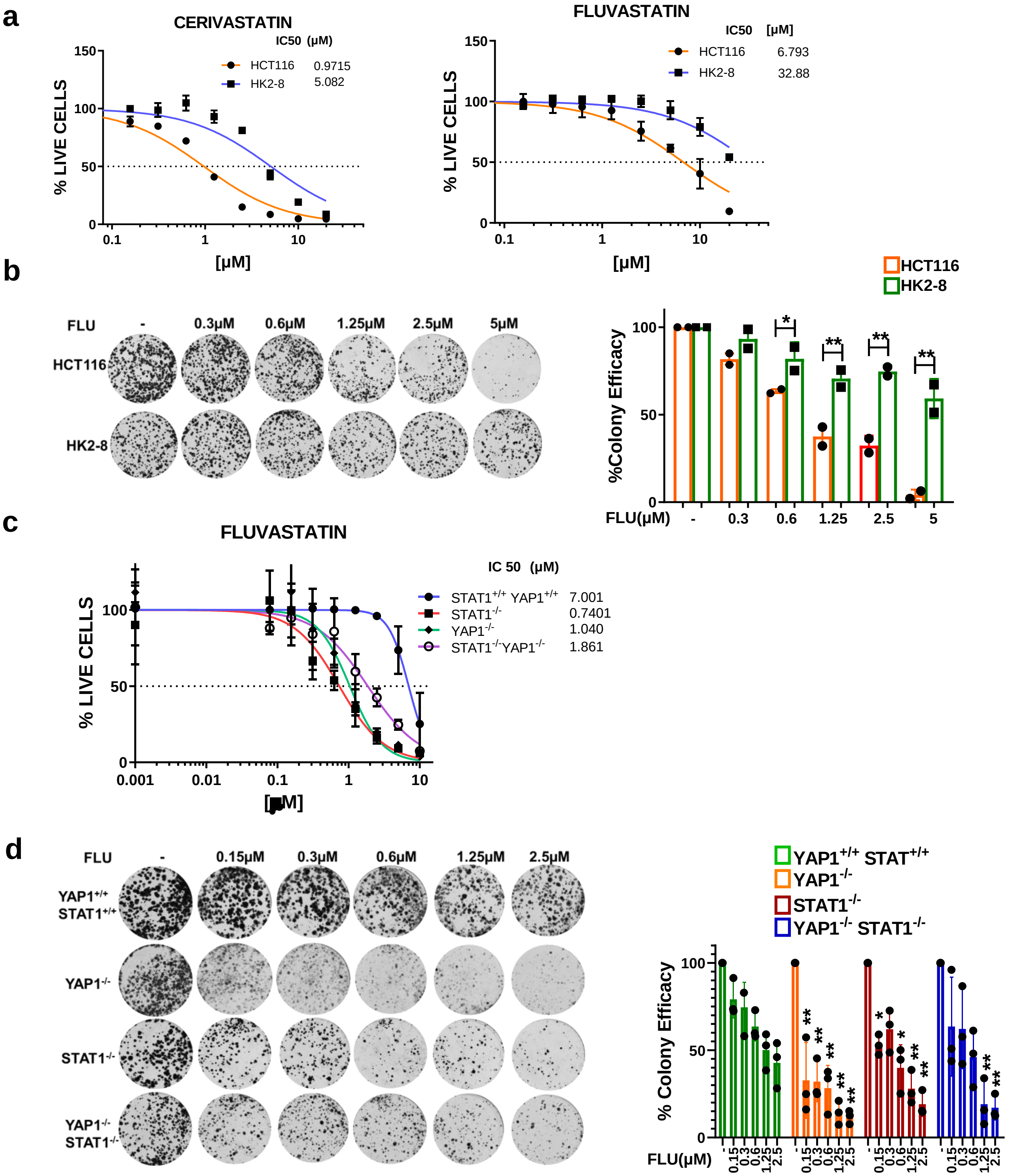

**Suppl Fig. 5.** ***STAT1 and YAP1 contribute to the resistance of mutant KRAS cells against inhibitors of the mevalonate pathway***. (**a**) $IC_{50}$ assays were conducted to compare the sensitivity of isogenic HCT116 and HK2-8 cells to treatment with cerivastatin or fluvastatin. (**b**) The colony-forming ability of HCT116 and HK2-8 cells was evaluated in response to increasing concentrations of fluvastatin. (**c**) $IC_{50}$ assays were also performed on HCT116 cells that were either replete or deplete for STAT1, YAP1, or both, following treatments with fluvastatin. (**d**) The colony-forming efficacy of HCT116 cells, replete or deplete for STAT1, YAP1, or both, was assessed in the presence of the indicated concentrations of fluvastatin. (**b**, **d**) Data were obtained from 3 independent experiments and represent ±SEM (*) $P < 0.05$; (**) $P < 0.01$; (*t*-test)

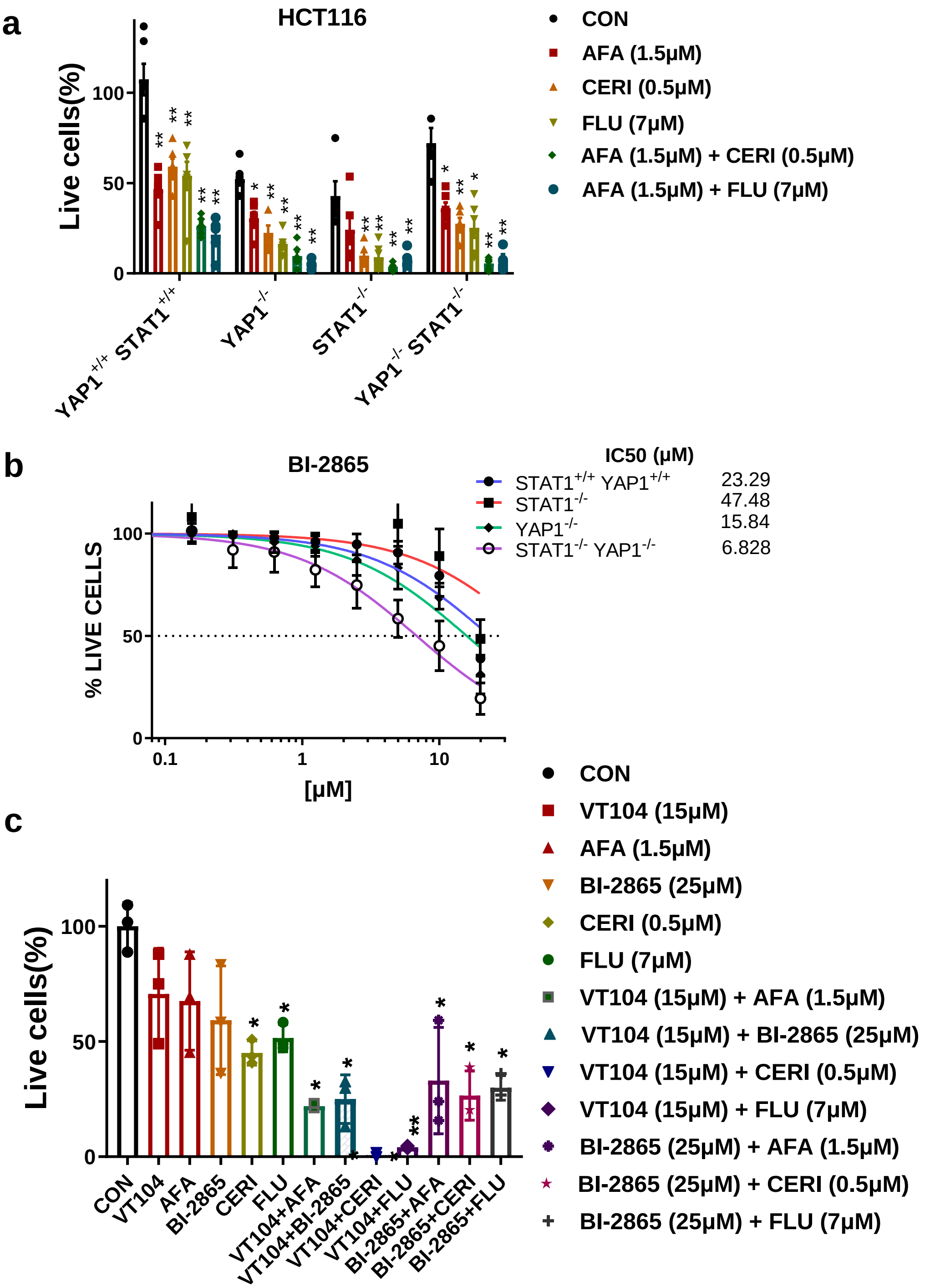

**Suppl Fig. 6.** ***The STAT1-YAP1 axis contributes to the resistance of mutant KRAS cells to chemotherapeutic drugs***. (**a**) The anti-proliferative effects of single and combination treatments using afatinib, fluvastatin, or cerivastatin were evaluated in HCT116 cells either containing or lacking STAT1, YAP1, or both. Cells were treated with the indicated concentrations of the drugs, which correspond to the $IC_{50}$ values determined from monotreatment assays. (**b**) $IC_{50}$ assays were performed on HCT116 cells that were either replete or deplete for STAT1, YAP1, or both, following treatments with BI-2865. (**c**) HCT116 cells were treated with single and combination therapies using the indicated anti-tumor drugs. The concentrations of afatinib, cerivastatin, fluvastatin, and BI-2865 used in the treatments corresponded to their respective $IC_{50}$ values, as determined in HCT116 cells. (**a**, **c**) Data were obtained from 3 independent experiments and represent ±SEM (*) $P < 0.05$; (**) $P < 0.01$ (*t*-test)

| | | | |
|---|---|---|---|
| **APPLICATION** | | **TARGET GENE** | **SEQUENCE** |
| **gRNA** | | YAP1 | #1: CAUCAGAUCGUGCACGUCCGCGG |
| | | | #2: AUCAGAUCGUGCACGUCCGCGGG |
| | | STAT1 | #1: GAGGUCAUGAAAACGGAUGGUGG |
| | | | #2: GCUUUUCUAACCACUGUGCCAGG |
| **siRNA** | | YAP1 | #1: GGUCAGAGAUACUUCUUAA |
| | | | #2: CCACCAAGCUAGAUAAAGA |
| | | | #3: GAACAUAGAAGGAGAGGAG |
| | | | #4: GCACCUAUCACUCUCGAGA |
| | | STAT1 | #1: AGAAAGAGCUUGACAGUAA |
| | | | #2: UAAAGGAACUGGAUAUAUC |
| | | | #3: GAGCUUCACUCCCUUAGUU |
| | | | #4: GAACCUGACUUCCAUGCGG |
| | | TEAD4 | #1: GACAGAGUAUGCUCGCUAU |
| | | | #2: UCAAGCACCUCCCUGAGAA |
| | | | #3: GGACACUACUCUUACCGCA |
| | | | #4: GUGCAUUGCCUAUGUCUUU |
| **qPCR** | | ACAT1 | Fw 5'-CAAGGCAGGCAGTATTGGGT-3' |
| | | | Rv 5'-TGGCTTTCATTCCTGAAGCAC-3' |
| | | HMGCR | Fw 5'-TGTCCAAAGTTTGAAGAGGATGT-3' |
| | | | Rv 5'-AGGATGGCTATGCATCGTGT-3' |
| | | IDI1 | Fw 5'-GCTTCTGCTACAGCAAAGATCAG-3' |
| | | | Rv 5'-AAGCTCGGCTGGATTGCTTA-3' |
| | | GAPDH | Fw 5'-CCTCCCGCTTCGCTCTCT-3' |
| | | | Rv 5'-CCGTTGACTCCGACCTTCAC-3' |
| | | ACTIN | Fw 5'-CAGCAGATGTGGATCAGCAAG-3' |
| | | | Rv 5'-GCATTTGCGGTGGACGAT-3' |
| **ChIP** | | SREBF1 STAT1 SITE #1 | Fw 5'-AAAAGCACACTCCCCACATC-3' |
| | | | Rv 5'- CTTGCCAAAAGCCAAAAGAG-3' |
| | | SREBF1 STAT1 SITE #2 | Fw 5'-AAGAGAATCCCCGAGGTGGT-3' |
| | | | Rv 5'-GGATGTGGGGAGTGTGCTTT-3' |
| | | SREBF2 STAT1 SITE #1 | Fw 5'- GTGTTGCCAGCACATCCAAG-3' |
| | | | Rv 5'-CCCATCCGTCCTAAGTCGAAG-3' |
| | | SREBF1 TEAD4 SITE | Fw 5'-GGACAGGAGTGGCCAAAGAA-3' |
| | | | Rv 5'-GCTTCCTCCTTCAGGAGGTG-3' |
| | | SREBF2 TEAD4 SITE | Fw 5'-AAGATGACGTAATGTGCTCCCA-3' |
| | | | Rv 5'-CACACGCAAGTCCGTCCA-3' |

**Supplementary Table 1.** Sequence of gRNA, siRNA and DNA primers used in the study.

| ANTIBODY | SPECIES | SOURCE | ID | | APPLICATION | |
|---|---|---|---|---|---|---|
| | | | | WB (xDILUTION) | IF (xDILUTION) | CHIP (xDILUTION) |
| STAT1α (C-111) | MOUSE MONOCLONAL | SANTA CRUZ | SC-417 | 1000 | | |
| STAT1(D1K9Y) | RABBIT MONOCLONAL | CELL SIGNALING TECHNOLOGY | 14994 | 1000 | | 50 |
| STAT1 pSer727 | RABBIT POLYCLONAL | CELL SIGNALING TECHNOLOGY | 9177 | 1000 | | |
| STAT3 (79D7) | RABBIT MONOCLONAL | CELL SIGNALING TECHNOLOGY | 4904 | | | 50 |
| STAT5(D2O6Y) | RABBIT MONOCLONAL | CELL SIGNALING TECHNOLOGY | 94205 | | | 50 |
| GFP | MOUSE MONOCLONAL | CELL SIGNALING TECHNOLOGY | 2955 | 1000 | | |
| SREBP1 | MOUSE MONOCLONAL | NOVUS BIO | NB600-582SS | 1000 | | |
| SREBP2 | RABBIT POLYCLONAL | NOVUS BIO | NB100-74543 | 1000 | | |
| HMGCR(A9) | MOUSE MONOCLONAL | DR. RUSSELL DEBOSE-BOYD | PMID: 22143767 | 1000 | | |
| YAP (D8H1X) XP | RABBIT POLYCLONAL | CELL SIGNALING TECHNOLOGY | 14074 | 1000 | 200 | |
| FAS(EPR7465) | RABBIT MONOCLONAL | ABCAM | AB128856 | 1000 | | |
| TEAD4/TEF-1 | RABBIT MONOCLONAL | ABCAM | AB197589 | 1000 | | |
| GFP-TRAP AGAROSE | VHH NANOBODY | CHROMO TEK | AB_2631357 | | | 50 |
| ACTIN | MOUSE MONOCLONAL | MB BIO | C4 | 5000 | | |
| THOC-1 (29) | MOUSE MONOCLONAL | SANTA CRUZ | SC-136426 | 1000 | | |
| TUBULIN | MOUSE MONOCLONAL | SIGMA-ALDRICH | T5168 | 1000 | | |
| MOUSE IgG-HRP | GOAT POLYCLONAL | KPL | 474-1806 | 3000 | | |
| RABBIT IgG-HRP | GOAT POLYCLONAL | JACKSON IMMUNE RESEARCH | 111-035-144 | 3000 | | |
| ALEXA FLUOR546 ANTI-RABBIT | GOAT POLYCLONAL | MOLECULAR PROBES | A11010 | | 2000 | |

**Supplementary Table 2.** Primary and secondary antibodies used in the study.
